# Molecular attributes of the tropical tree *Avicennia schaueriana* involved in the response and tolerance to low temperatures

**DOI:** 10.1101/2024.02.08.579386

**Authors:** Yohans Alves de Moura, Alexandre Hild Aono, Mariana Vargas Cruz, Alessandro Alves Pereira, João de Deus Vidal, Anete Pereira de Souza

## Abstract

Mangroves are coastal ecosystems of great socioenvironmental importance that are highly threatened by human activities. Mangrove trees live under harsh environmental conditions, which makes them sensitive to extreme weather events, particularly freezing events. Such events are unpredictable and have catastrophic consequences for mangrove trees; therefore, understanding and anticipating the impacts of such events are essential for directing future mitigation measures. Freezing cold currently limits the distribution of mangroves to tropical and subtropical latitudes worldwide. Mangrove trees are seriously affected by freezing conditions and suffer severe metabolic fluctuations due to photosystem and cellular structure damage. However, land plants more broadly have developed sophisticated mechanisms of resistance to freezing during their evolution, and the central molecular mechanisms involved in this process are consistent. However, the known information is restricted to models of herbaceous plants, such as *Arabidopsis thaliana*, that are native to temperate habitats, and there is a research gap regarding tropical trees such as mangroves. This work aimed to improve the understanding of the molecular aspects of the response and tolerance to freezing in mangrove trees using *Avicennia schaueriana* as a model. This species occurs within the colder range limits of South American mangroves and shows evidence of the existence of two functional groups that are locally adapted to the equatorial (EQ) and subtropical (ST) portions of the Brazilian coast. We investigated the transcriptional profiles of seedlings from both functional groups under freezing shock (−4°C) in a time series. We analyzed transcriptomic data by combining differential expression, coexpression network and protein interaction data. Our results allowed us to describe the profile of the molecular response of *A. schaueriana* to freezing and the divergence in the behavior of the EQ and ST functional groups. In EQ plants, the response strongly depended on the action of abscisic acid (ABA) and stress signals throughout the experiment. Notably, ABA negatively affects plant growth and promotes the accumulation of carotenoids, anthocyanins and lipids through chlorophyll degradation. On the other hand, in the ST, there were fewer hormones active in the process of primary growth maintenance and metabolic normalization. The accumulation of substances is mainly based on sucrose, anthocyanin and lipid levels, and lipid synthesis is not dependent on chlorophyll degradation. Based on these results, we hypothesize that susceptibility to freezing damage is greater in EQ mangroves than in ST mangroves. Therefore, we recommend that this fact be considered when managing this species, especially at higher latitudes, which are more prone to lower temperatures and extreme freezing events.

## INTRODUCTION

Mangroves are coastal ecosystems of great global socioenvironmental importance that enhance climate regulation, maintenance of marine diversity and physical protection of the coast (Eong 1993, Sandilyan & Kathiresan 2012, Vo et al. 2012). Despite their importance, they are seriously threatened by human activities such as deforestation and climate change (Hamilton & Friess 2018). The mangrove ecosystem has a delicate balance because its organisms, especially trees, live under severe abiotic conditions involving high rates of salinity, insolation and flooding (Bajay et al. 2018 and Curz et al. 2019). This delicate balance makes them particularly sensitive to the occurrence of extreme climatic events (ECEs), especially freezing events (Chen et al. 2010). Moreover, such events are increasing in frequency and intensity, are unpredictable and have catastrophic environmental consequences (Ummenhofer et al. 2017). Therefore, understanding and anticipating the consequences of ECEs are extremely important for future mitigation of their impacts.

Low temperatures (< 15°C) and freezing events (< 0°C) are the primary limitations for the geographic range of mangroves, with these events historically shaping mangrove distributions and currently limiting them to tropical and subtropical latitudes worldwide (Tomlinson 1986, Coock-Patton et al. 2015 and Osland et al. 2017). Mangrove trees are the structural element of this ecosystem and experience cold as an especially severe stressor (Ritonga & Chen 2020). Indeed, plant metabolism is strongly impacted at the cellular and systemic levels during cold events (Deng et al. 2019). Among the most striking impacts are the risk of cavitation and xylem embolism (Charreir et al. 2014), reductions in metabolism, and restrictions in photosynthetic capacity (Chinnunsamy et al. 2007, Sevillano et al. 2009). These processes culminate in high levels of reactive oxygen species (ROS) and loss of cell membrane fluidity (Tyystjärvi 2013, Ding et al. 2019). When this occurs, the central processes for plant survival are significantly affected, limiting growth and increasing the risk of death from desiccation and cell collapse (Ding et al. 2019).

It is theorized that land plants have developed a series of sophisticated mechanisms to resist low temperatures and freezing during their evolution, allowing occupation of colder environments (Sinnot and Bailey 1915, Wing and Boucher 1998, Vermeji and Herbert 2004, Ding et al. 2019, Ritonga & Chen 2020). These mechanisms include important morphophysiological adaptations, such as deciduous leaves, herbaceous habits and narrower and more resistant vascular elements (Zane et al. 2014). The accumulation of substances such as soluble sugars, lipids, amino acids, pigments, and other osmolytes also helps to stabilize plant cellular structures during cold and freezing events (Nishida & Murata 1996, Zhang et al. 2016, Deng et al. 2019 and Bilska-kos et al. 2020).

Plant physiological and molecular responses that confer cold and freezing tolerance require intensive reprogramming of gene expression and metabolism (Ydav 2010). The molecular mechanisms involved in the regulation of phenotypic and physiological changes in plants in response to freezing conditions are currently being elucidated due to advancements in massive genetic sequencing techniques. For example, the CBF (C-repeat/DRE-binding factor) gene family has been the main target of research in the last 20 years (Ding et al. 2019, Ritonga & Chen 2020). These genes are part of the APETALA2/ETHYLENE RESPONSE FACTOR (AP2/ERF) superfamily and play a central role in inducing the expression of genes linked to the cold response, COR, in *A. thaliana* (Medina et al. 1999, Fowler & Tomashow 2002, Jia et al., 2016). The expression of COR genes directly results in metabolic responses, such as the accumulation of antioxidant substances and osmolytes, which confer greater tolerance to cold and freezing (Ritonga & Chen 2020).

The expression of CBF-COR and other cold tolerance pathways can be influenced by phytohormones such as abscisic acid (ABA) and ethylene (ETH) and through posttranslational protein changes such as phosphorylation/dephosphorylation and ubiquitination (Eremina et al. 2016, Ding et al. 2019). Hormones such as ABA and ETH can directly induce the expression of CBF-COR, the same pathway that induces desiccation tolerance (Eremina et al. 2016). Posttranslational phenomena can affect hormone signal propagation pathways in many ways. For example, phosphorylation events are necessary for the propagation of ABA and ETH signals (Raz & Fluhr 1993, Yang et al. 2017). On the other hand, ubiquitination negatively impacts ABA synthesis and signaling, modulating these processes at many steps during stressful events (Xu & Xue 2019). Therefore, the metabolic reprogramming linked to the response to cold and freezing involves various complex elements, and additional studies are needed to better understand this topic.

Despite recent advances in understanding the central molecular mechanisms involved in tolerance of low temperatures, most of this research has focused on herbaceous plants, such as the model species *A. thaliana,* and some domesticated plants, such as tomato (Chen et al. 2015) and rice (Min et al. 2016). This topic has rarely been explored in tropical trees such as mangroves, except for in the work of Fei et al. (2015, 2021), who investigated a few genes active in *Kandelia obovata*, an endemic species of Southeast Asia (Duke 2017). Studies involving trees are often limited by difficulties in implementing traditional research methodologies. These difficulties are due to the long lifespan of trees combined with the lack of prior biological and genomic information related to them (Holliday et al. 2017). Importantly, improving the understanding of the molecular mechanisms underlying tolerance to low temperatures in mangroves can aid in their conservation and increase the understanding of the evolutionary ecology of trees in cold environments.

Currently, the use of the RNA-seq technique (Wetterstrand 2018) and its combination with statistical methods such as de novo assembly of transcriptomes, differential expression analyses and functional enrichment have proven to be effective ways to overcome the lack of previous genomic information (Wu et al. 2012, Van De Wiel et al. 2013, Love et al. 2014, Bajay et al. 2018). Coexpression networks have also helped in this respect, allowing inference of the biological and regulatory functions of genes and groups of genes with unknown activity (Rao & Dixon 2019). Additionally, common garden experiments have been applied together with new transcriptomic techniques. This type of experiment allows direct control of the exposure of plants from different geographic origins to stressors and has been used successfully for studies related to molecular responses in plants under abiotic stresses (O’Rourke et al. 2013, Wang et al. 2014, Akman et al. 2016).

In this scenario, mangroves of the genus *Avicennia* emerge as an interesting model for studying the impacts of cold on mangroves. *Avicennia* trees are the most resistant mangroves to low temperatures and occur in all tropical and temperate mangroves in the world, such as *A. germinans* on the coast of the USA and *A. marina* in South Africa and Australia (Morrisey et al. 2010). In South America, most species occur along the Brazilian coast, with *A. schaueriana* being the most representative and ubiquitous species along the entire coast of the country except in the extreme southern region. In Brazil, this species lives from the equatorial Amazon region to the latitudinal limit of South American mangroves at 28°3’S and resists the lowest temperatures tolerated by the ecosystem on the continent (Soares et al. 2012). This broad latitudinal gradient causes the species to experience varied environmental conditions, with allelic and transcriptional evidence for the existence of two locally adapted groups: the equatorial group (EQ) and the subtropical group (ST) (Figure 1) (Cruz et al. 2019). The EQ group occurs in environments with more stable, hot and dry conditions, while the ST group occurs in seasonal, humid and cold environments. Given these characteristics, we selected *A. schaueriana* as a model for the molecular study of South American mangroves under freezing temperatures.

**Figure 1:**
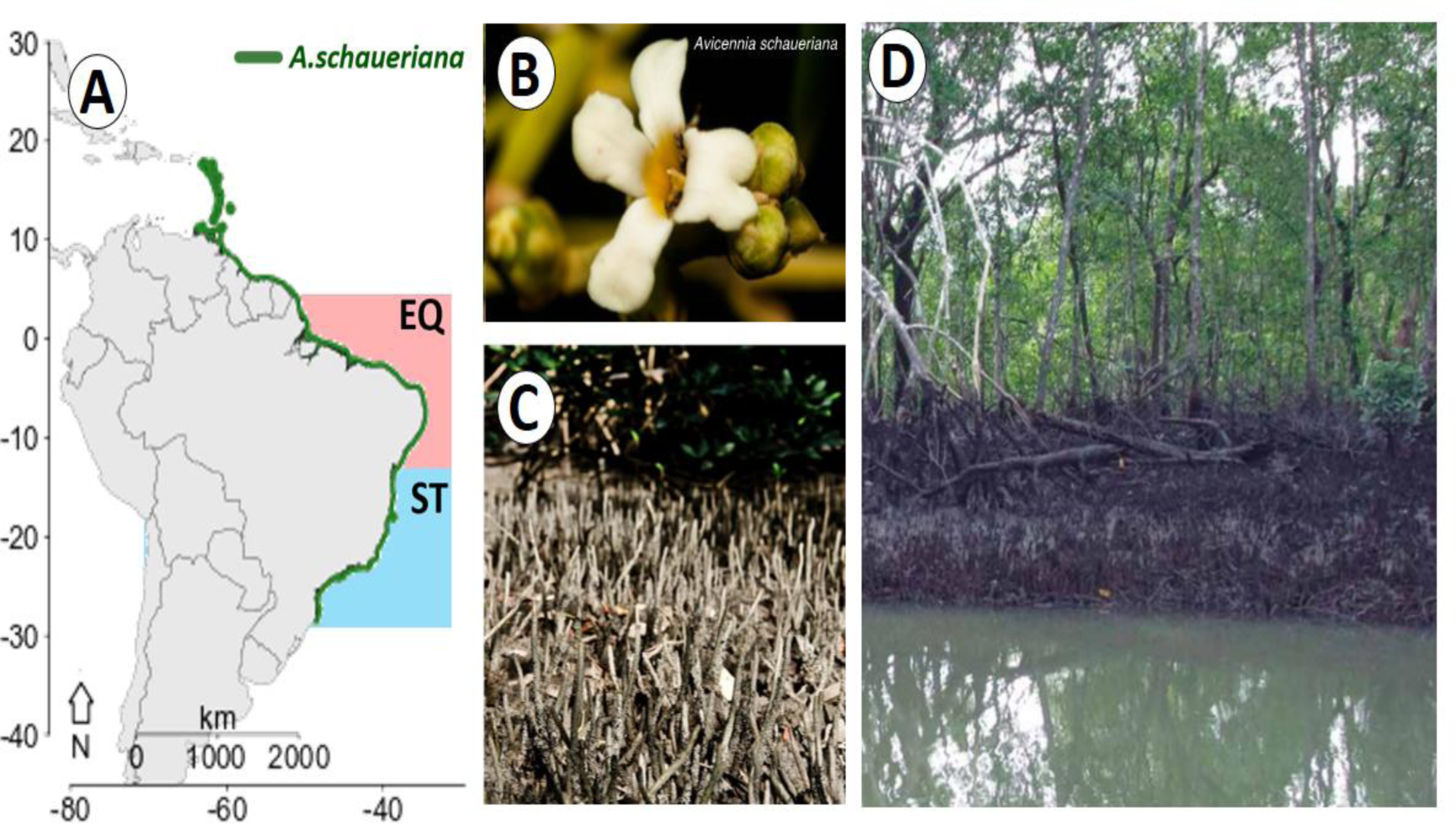
(A) Distribution of *Avicennia schaueriana* along the South American Atlantic coast and the delimitation of the EQ and ST functional groups, (B) detail of the *A. schaueriana* flower, (C) detail of the pneumatophore roots characteristic of *Avicennia*, and (D) view of *A. schaueriana* trees. Sources: (B) and (C) Gustavo M. Mori and (D) Atlas of Brazilian mangroves – MMA/ICMBIO 2018.

In this work, we aimed to add to the body of knowledge on the molecular aspects involved in the response and tolerance to freezing in mangrove trees. To this end, we investigated the transcriptional profile of *A. schaueriana* seedlings from EQ and ST latitudes on the Brazilian coast under freezing shock, thus determining the functional diversity of this species. The plants were grown under homogeneous environmental conditions in a common garden to reduce potential confounding factors from both functional groups linked to different local climates. The analysis of expression levels was conducted by combining differential expression techniques and transcript and protein coexpression networks. In the latter, we investigated the EQ and ST groups separately to identify transcripts and proteins that were possibly linked to regulatory functions during the experiment. Our results provide important information on this topic and valuable insights for future mangrove management, conservation and restoration programs on the South American Atlantic coast. Overall, we hypothesized that, while still under stress, ST plants would exhibit greater resistance to freezing due to signs of continued growth, together with more sustainable lipid synthesis and metabolic recovery. Our findings showed that, when subjected to freezing conditions, the EQ plants exhibited signs of growth inhibition, degradation of chlorophyll as a lipid source and permanent stress-related metabolic signs.

## MATERIALS AND METHODS

### Common garden experiment - freezing shock

Mature *A. schaueriana* propagules were collected from five mother trees at least 100 m apart in two populations, i.e., the EQ and ST groups. Plants in these groups are present at contrasting latitudes and are therefore subject to contrasting temperature regimes. For the ST group, the chosen location corresponds to the southern limit of mangroves in the Americas, which is in a subtropical region (ST) of the state of Santa Catarina (Soares et al. 2012) (Figure 2). For the EQ group, the location corresponds to the largest continuous belt of mangrove forest in the world, and the Macromarés Mangrove Coast of the Amazon is located in the state of Pará (Kjerfve et al. 2002, Souza-Filho et al. 2006) (Figure 2).

**Figure 2:**
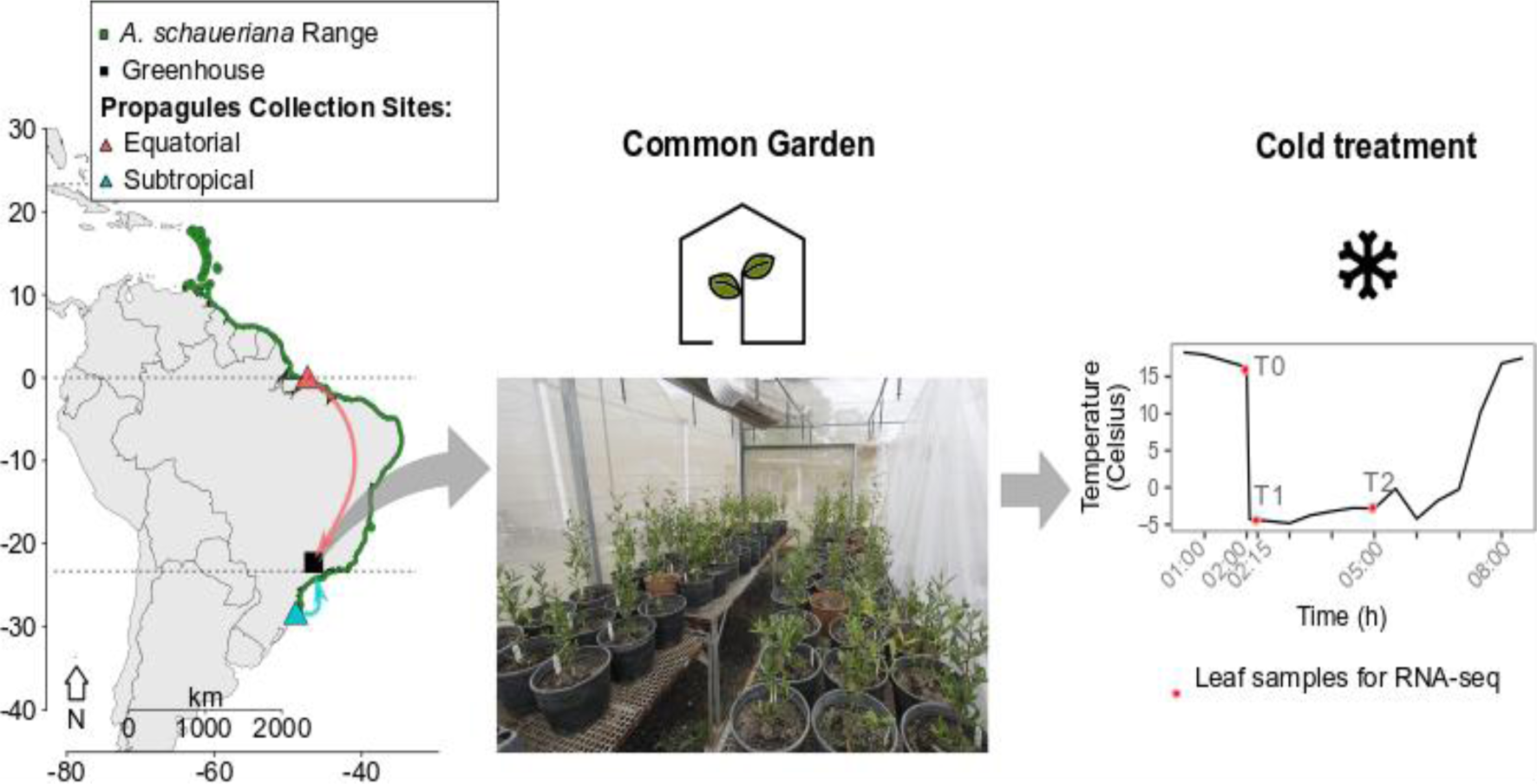
General scheme of the common garden experiment. Propagule collection sites: Growth under homogeneous conditions in a greenhouse and subsequent exposure to low temperature (average of −4.0°C). The red dots on the graph represent the three sampling time points (T0, T1 and T2).

The propagules were germinated in mangrove soil from their respective places of origin following the protocol described by Reef and colleagues (Reef et al. 2015) for *Avicennia germinans* and subsequently grown in trays for two months. After this period, the plants were transferred to 6-liter pots with sand and topsoil (1:1), where they were grown for seven months in a greenhouse under homogeneous conditions at the State University of Campinas (UNICAMP) (22°49’ S 47°04’ W). The plants were watered twice a week with 0.4× Hoagland solution plus 15 g/L NaCl. Pot rotation was carried out weekly to avoid the effects of environmental heterogeneity on plant growth.

Five seedlings from different mother trees from each of the two locations (equatorial and subtropical) were used to test the response to freezing. All ten plants were exposed to a temperature of −4°C, and leaf material was collected before the start of treatment (T0), 15 minutes after the start of treatment (T1) and 180 minutes after the start of treatment (T2) (Figure 2). Young leaves were collected for RNA extraction. The time intervals used were adapted from the 2011 study carried out by Carvallo and collaborators with *A. thaliana*, in which 15 minutes was the duration at which transcription factor expression was responsive to cold and 3 hours was the duration at which genes affected by transcription factors were expressed.

### RNA extraction and sequencing

RNA was extracted from leaves sampled according to the protocol described by Oliveira et al. (2015), with three sampling time points (T0, T1 and T2) for each of the five plants from the two different locations (EQ and ST). The integrity and purity of these samples were evaluated on a 1% agarose gel and on a NanoVue spectrophotometer (GE Healthcare Life Sciences, Chicago IL, US). We used an Illumina paired-end TruSeq RNA library preparation kit for 150 sequencing cycles for a mid-output flowcell. Each sample was marked with specific barcodes that made it possible to identify the origin of the sequences obtained at the end of the sequencing process. The libraries were subsequently sequenced on an Illumina NextSeq 500 platform (Illumina, Inc.).

### “De novo” assembly and transcriptome characterization

Sequence quality was checked with *FastQC* software (Andrews 2010), and filtering was performed with *Trimmomatic 0.36* software (Bolger et al. 2014). De novo transcriptome assembly was performed using *Trinity 2.5.1* software (Grabherr et al. 2011) with a minimum length of assembled transcripts of 200 bases and a tenfold minimum depth per base. Verification of the structural quality of the assembly as well as the unigene version of the transcriptome was performed using *Python* and *Perl* scripts contained in the *Trinity 2.5.1* package (Grabherr et al. 2011). Transcriptome completeness was analyzed via *BUSCO 3.1.10* (Waterhouse et al. 2018), in which assembled unigene sequences were aligned against universal ortholog sequences from all land plants (Viridiplantae).

The prediction of potentially coding regions was performed with *Transdecoder v5.5* software (Haas & Papanicolau 2017). In this step, start and stop codons were identified in the DNA sequences and used to determine coding intervals for proteins or parts thereof. For functional annotation, *Trinotate v3.2.1* software (Bryant et al. 2017) was used, and the predicted gene sequences were compared via BLAST 2.5.1 (Camacho et al. 2009) against the UniProt-SwissProt, TAIR and NR-NCBI databases. Protein domains were identified via HMMER against the Pfam database (Finn et al. 2014), and gene ontology (GO) terms were obtained from the UniProt-SwissProt and TAIR and NR databases following the *Trinotate v3* software pipeline .2.1 (Bryant et al. 2017).

### (I) Identification of differentially expressed transcripts

Mapping and counting of sequences aligned against the unigene version of the assembled transcriptome was performed with *Salmon 0.5* software (Patro et al. 2017). Differentially expressed transcripts were identified using the *edgeR* (Robinson et al. 2010) and *lima-voom* (Ritchie et al. 2015) packages in R 4.0.3 (R Core Team 2020). Transcripts with fewer than ten counts were removed, and subsequently, the counts were normalized with edgeR (Robinson et al. 2010) using the TPM method (transcripts per million kilobase). The normalized values were converted to a base 2 logarithmic scale (log 2 FC); only transcripts with log2FC > |1| and a false positive discovery rate (FDR) < 0.05 (obtained by Bonferroni correction) were retained for further analyses.

Initially, an exploratory multidimensional scaling analysis was carried out with the *edgeR* package. At this stage, the similarity of the samples according to the expression profile was verified. Subsequently, comparisons were made between individuals from two different populations, EQ (equatorial) and ST (subtropical), at three time points after exposure to cold (−4°C), with T0 being the control, T1 = 15 minutes and T2 = 180 minutes. Comparisons are made *within the time points* EQ_T0 × ST_T0; EQ_T1 × ST_T1; EQ_T2 × ST_T2; and *within each population over time,* EQ_T0 × EQ_T1; EQ_T0 × EQ_T2; EQ_T1 × EQ_T2; ST_T0 × ST_T1; ST_T0 × ST_T2; and ST_T1 × ST_T2.

### DET functional enrichment–gene ontology (GO) analysis

Enrichment analysis of gene ontology (GO) terms associated with differentially expressed transcripts (DETs) was performed with the *TopGo* package (Alexa and Rahnenfuhrer 2020) in R 4.0.3 (R Core Team 2020) using the weight01 algorithm. This algorithm considers the hierarchy levels of GO terms and minimizes losses in the identification of false positives based on the Fischer test. The p value used was 0.01, and the test was performed for all comparisons mentioned in the previous subsection. All GO terms with statistically significant enrichment for the biological processes (BP) domain were retained.

### DET functional enrichment–enzymatic pathways (KEGG)

Each of the DETs has been associated with a single KEGG pathway (Kanehisa & Goto 2000). Only pathways with corresponding annotations in *A. thaliana* were considered to minimize interpretation errors. The enriched pathways were detected by Fisher’s test performed using R software (R Core Team 2020), and only pathways with a p value less than 0.05 were retained.

### (II) Coexpression network analyses Global coexpression network and functional gene groups

To understand the synergistic effects of the transcripts, we used a network construction method based on weighted gene coexpression network analysis (WGCNA) (Langfelder & Horvath 2008). Briefly, this method maximizes the strongest connections between transcripts and maintains them at the expense of the weakest ones, thus allowing the generated networks to have a topology as close as possible to a scale-free network. The absence of scales is an important property for the study of nonrandom and complex processes such as gene interactions. This property guarantees that genes absent in networks, motivated by limitations in sequencing or assembly, do not alter other correlations (Barbasi & Oltvai 2004). In this way, scale-free networks consist of a statistically robust system for using sample data to estimate real values.

The networks were constructed using the *WGCNA* package (Langfelder & Horvath 2008) in R 4.0.3 (R Core Team 2020). We used the normalized expression values (TPMs) to construct a global coexpression matrix based on the Pearson correlation index (Pearson 1986). The correlation values were subsequently transformed and weighted by testing several different powers (powers β) to detect networks with scale-free topologies. The groups of transcripts with the highest correlation among themselves compared to the others were subsequently ordered into functional groups, also called modules.

### DET-enriched modules - Identification and temporal categorization

Modules enriched with differentially expressed transcripts (DETs) were identified via Fisher’s test using R 4.0.3 software (R Core Team 2020), and only those with p values < 0.05 were retained. Subsequently, the GO terms associated with biological processes enriched in these modules were identified with the *TopGo* package (Alexa and Rahnenfuhrer 2020) in R 4.0.3 (R Core Team 2020) using the weight01 algorithm, with only terms related to biological processes with p values < 0.01.

After identifying the enriched modules, the temporal behavior of enrichment in DETs in each of the EQ and ST functional groups was verified. The modules were categorized into six temporal classes, the first reflecting the constancy of enrichment throughout the experiment (T0, T1 and T2), called *basal processes*. The second and third modules were enriched at T1 and T2, respectively; these modules are called *fast activation* and *slow activation*, respectively. The fourth and fifth categories contained the modules whose enrichment in DETs stopped at T1 and T2, respectively, called *fast suppression* and *slow suppression*. Finally, the sixth category included modules whose enrichment in DETs stopped at T1 and was restored after T2; these were called *suppression* and *restitution*.

Initially, the functional enrichment results for each module were analyzed individually, and modules with highlighted functionality under freezing conditions were selected for each temporal class and functional group. These modules were used in the next step to construct specific networks for EQ and ST. Subsequently, the GO terms referring to the biological processes of each temporal class were summarized with the *REVIGO* algorithm (Supek et al. 2011). The criterion used was that terms with higher absolute numbers of repetitions be more representative, with an average redundancy cutoff of 0.7. The summarized terms and their ontogenetic relationships were graphically represented by Cytoscape 3.9.1 software (Shannon et al. 2003).

### Specific population network and hub identification

The group of modules with highlighted importance under freezing conditions was used as a basis for building new coexpression networks. Only information relating to the expression levels of transcripts contained in these modules was used to construct specific networks, one for the EQ group and the other for the ST group. The networks were constructed using the *Algorithm for the Reconstruction of Accurate Cellular Networks (ARACNE)* method executed with the *MINET* package (Meyer et al. 2008) in R 4.0.3 software (R Core Team 2020).

In the subsequent step, we focused our topological analysis on identifying the hub transcripts in each specific network. Biologically, the expression of these transcripts is connected in a very consistent way to the other transcripts and can strongly impact general expression patterns. This property allowed us to infer possible regulatory functions of the hubs. Mathematically, hubs are the nodes that have the greatest number of connections with other nodes in the network. In this way, the probabilities of each transcript concentrating on the connections between the expression patterns were calculated using *Kleimberg’s Hubscore* statistic (Kleimberg 1999) in the software R 4.0.3 (R Core Team 2020). We retained only the transcripts with the five highest values from each network. After identifying the likely hubs, we used the annotation information at the transcript level to interpret the results.

### Protein–protein interaction network

The protein interaction networks were constructed from the transcripts contained in the cold-sensitive modules identified in the previous steps. These transcripts were associated with *A. thaliana* pathways related to proteins and metabolic interactions. Data regarding interactions are available on the *STRING* website (Szklarczyk et al. 2023). The initial network was constructed with information from this group of transcripts. Subsequently, proteins that underwent immediate interaction with proteins present in the initial network (the 1st and 2nd neighbors) were also included. All manipulations and graphical visualizations of the network were performed using *Cytoscape* software (Shannon et al. 2003). After completing the network construction, the five proteins with the highest number of connections were identified as possible hubs.

### Differential expression validation

Nine differentially expressed transcripts were selected for RNA-seq validation through quantification by real-time amplification (RT–qPCR). The selection criteria were based on high LFC values and low p values, in addition to high absolute expression values. Two transcripts with similar and constant expression were used as normalizers and were selected based on the data from the experiment itself. The sequences of the primers used were designed with *Primer3Plus* software (Untergasser et al. 2007), limiting the length of the primers and amplicons to 20 and 120 bases, respectively.

All the RNA samples used in the experiment were amplified. Initially, we transformed 1000 ng of RNA into cDNA with the QuantTect Quiagem Reverse Transcription Kit. The reactions were subsequently performed in triplicate for each cDNA sample as follows: Sybr 2X, 4.2 μl; H2O Millipore, 1.4 μl; forward primer, 0.3 μl; reverse primer, 0.3 μl; and cDNA, 0.2 μl. The amplification program consisted of 7 steps: (1) 95°C for 3 minutes; (2) 95°C for 20 s; (3) 60°C for 20 s; (4) 72°C for 30 s; (5) repeated steps 2 to 4 39 time points; (6) 65°C for 5 s; and (7) 95°C for 5 s. The reactions were carried out in a CFX384 thermocycler Real Time System (Bio-Rad).

At the end of the reaction, amplification cycle counts (Cq) were used to measure the number of copies of each transcript present in each sample (expression levels). The relationship between Cq and the expression level is inversely proportional: the highest expression levels present amplifications in a smaller number of cycles due to the greater initial quantity of copies. The relative concentration (RQ) of each sample was determined using the formula RQ = - 2^ΔΔCq. After calculating the RQ, a t test was performed for all samples, retracing the contrasts made in the differential expression stage. Relative expression plots for each transcript selected in the contrast where it was detected as DET are shown in Supplementary Figure F16.

## RESULTS

### Transcriptome assembly and characterization

Sequencing was performed on Illumina NextSeq 500 platform, resulting in 95 million pairs of 75-base sense and antisense sequences. After filtering and quality selection, 67.85 million sequences were used to assemble the transcriptome. The assembled transcriptome contained 106,338 unigene transcripts, for which the N50 value was 1822 bases (Supplementary Table S1). The unigene transcriptome contains only the longest isoform of each assembled transcript and was used for all subsequent steps. The percentage of sequence pairs that were aligned to unigenes was 89%. BUSCO analysis revealed 84% similarity of the assembled transcripts against universal orthologs found in land plants (Supplementary Table S2).

Structural prediction revealed 52553 potential coding regions (ORFs) distributed in 31541 (29.6%) of the assembled transcripts. A total of 74847 (30.7%) transcripts did not contain potential coding regions. Functional annotation revealed 45% of the transcripts annotated from NCBI-NR, 31% from UniProt and 35% from TAIR.

### (I) Identification of differentially expressed transcripts (DETs)

During multidimensional scaling, samples were grouped by expression patterns with respect to the sampling groups, and the EQ and ST samples were grouped together. In temporal contrasts, the trend was repeated, and samples from similar time points emerged as discrete groups (Supplementary Figures F1 and F2). According to the differential expression test, 1432, 1605 and 1724 DETs were identified in the contrasts *between each functional group* (EQ and ST) at different time points (T0, T1, T2) (Figure 3a). Contrasts made *within each functional group* over time did not show statistically supported DETs.

**Figure 3:**
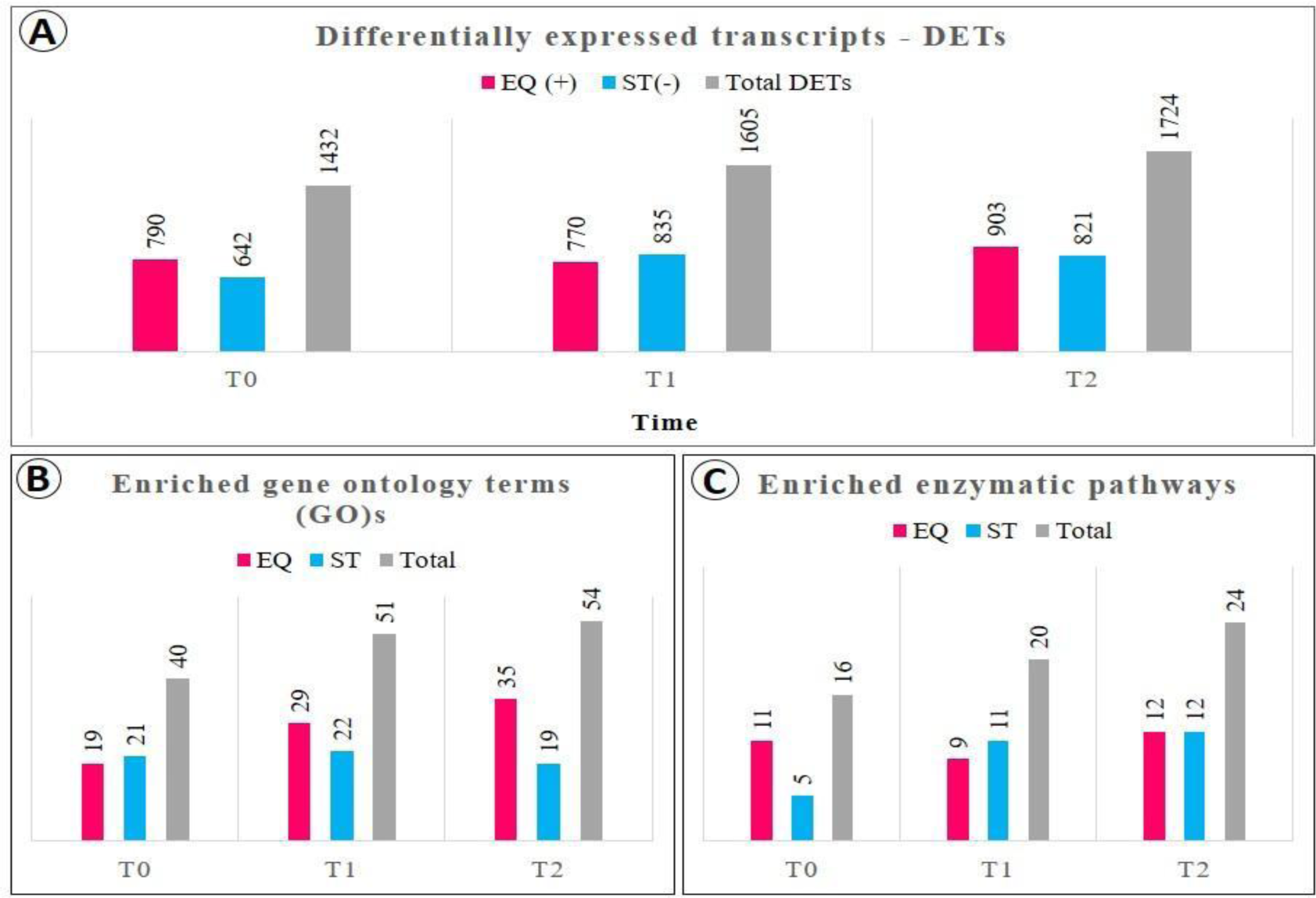
(A) Differentially expressed transcripts (DETs) obtained from comparisons between the equatorial (EQ) and subtropical (ST) groups of *A. schaueriana* exposed to −4°C at sampling time points T0, T1 (15 minutes) and T2 (180 minutes), with log2FC values > |1| and FDR < 0.05. (B) Numbers of GO terms associated with enriched biological processes among the DETs. (C) Enzymatic pathways enriched among DETs.

### DET functional enrichment–gene ontology (GO) analysis

The GO term enrichment test was performed only for comparisons between the EQ and ST groups within each of the T0, T1 and T2 time points, as we did not obtain differentially expressed transcripts in the comparisons *within each group over time*. We identified 242857 GO terms associated with 32854 transcripts in the transcriptome. Of these, 40 terms related to biological processes were significantly enriched at T0, 51 at T1 and 54 at T2. In general, the EQ group presented more enriched terms after exposure to freezing (T1 and T2) (Figure 5b).

### T0 - Control

The GO terms associated with biological processes that were linked to the development of reproductive structures, interaction with pathogens and the redox process were determined for both the EQ and ST groups. Specifically, in the EQ group, GO terms linked to phosphate ion transport, circadian regulation/photoperiodism, and abscisic acid (ABA) suppression were enriched. On the other hand, in the ST, biological processes linked to vacuolar salt and zinc sequestration, secondary metabolism and response to alkaloids were enriched (Supplementary Tables S3 and S4).

### T1-15 minutes after freezing

After the first time interval, biological processes linked to the development of reproductive structures, interactions with pathogens and redox reactions remained enriched in both groups. Specifically, in the EQ group, biological processes linked to the suppression of growth and transcription, response to drought, synthesis of alkaloids and carotenoids and synthesis of hormones (ABA, AS, GA and ETH) became enriched at T1. In the ST group, enrichment of genes was linked to the maintenance of primary growth, changes in DNA and the response to mannitol (Supplementary Tables S5 and S6).

### T2-180 minutes after freezing

After 180 minutes of freezing, biological processes linked to the development of reproductive structures, interactions against pathogens and oxidation‒reduction reactions remained enriched in both groups. Specifically, in EQ, growth suppression was enriched and was increased by amino acid synthesis, the anoxia response, hormone signaling (abscisic acid, ABA; auxins, AIA; ethylene, ETH; and cytokinins, CK) and protein ubiquitination. In the ST, the enriched processes were linked to cell wall reinforcement, the synthesis of strigolactones and anthocyanins, induced resistance via jasmonic acid (JA) and ETH, and protein dephosphorylation and transposition via RNA (Supplementary Tables S7 and S8).

### DET functional enrichment – enzymatic pathways (KEGG)

In total, 60 enriched pathways were found among all the contrasts (16, 20 and 24 at T0, T1 and T2, respectively). The number of enriched pathways per group was very similar, except at time point T0 (Figure 5c).

### T0 - control

At the control time point, the populations presented very different enriched pathway profiles. In the EQ group, the enriched pathways were linked to the synthesis of secondary compounds such as terpenoids and carotenoids, circadian regulation, interaction with pathogens and inositol phosphate metabolism. Pathways linked to glycerophospholipid metabolism, oxidative phosphorylation and hormonal signal transduction were enriched in the STs (Supplementary Table S9).

### T1-15 minutes after freezing

After 15 minutes of freezing in the EQ group, the pathways from the previous time point remained enriched, as did the flavonoid, tyrosine and ribosome synthesis pathways. In STs, the pathways from the previous period also remained enriched, as did the synthesis of more lipid types (sphingolipids and glycerolipids), galactose metabolism and ABC transporters (Supplementary Table S10).

### T2-180 minutes after freezing

After 180 minutes of freezing, in both groups, the pathways enriched at the previous time points were maintained. In the EQ treatment, the activity of the anthocyanin synthesis pathway increased. In the ST treatment, there was an increase in the pathways associated with brassinosteroid synthesis and sucrose, starch and inositol phosphate metabolism (Supplementary Table S11).

### (II) Coexpression network analysis Temporal categorization of the global network and modules

In the global network, 220 functional modules were identified, each containing 50 to 1228 transcripts, 64 of which were enriched with DETs. The *basal process* class presented the highest number of modules (34 in both the EQ and ST groups), while the classes affected by the freezing cold condition totaled 29 modules (Table 1). The EQ group presented a slightly greater number of enriched modules than did the ST group (Table 1).

**Table 1.**
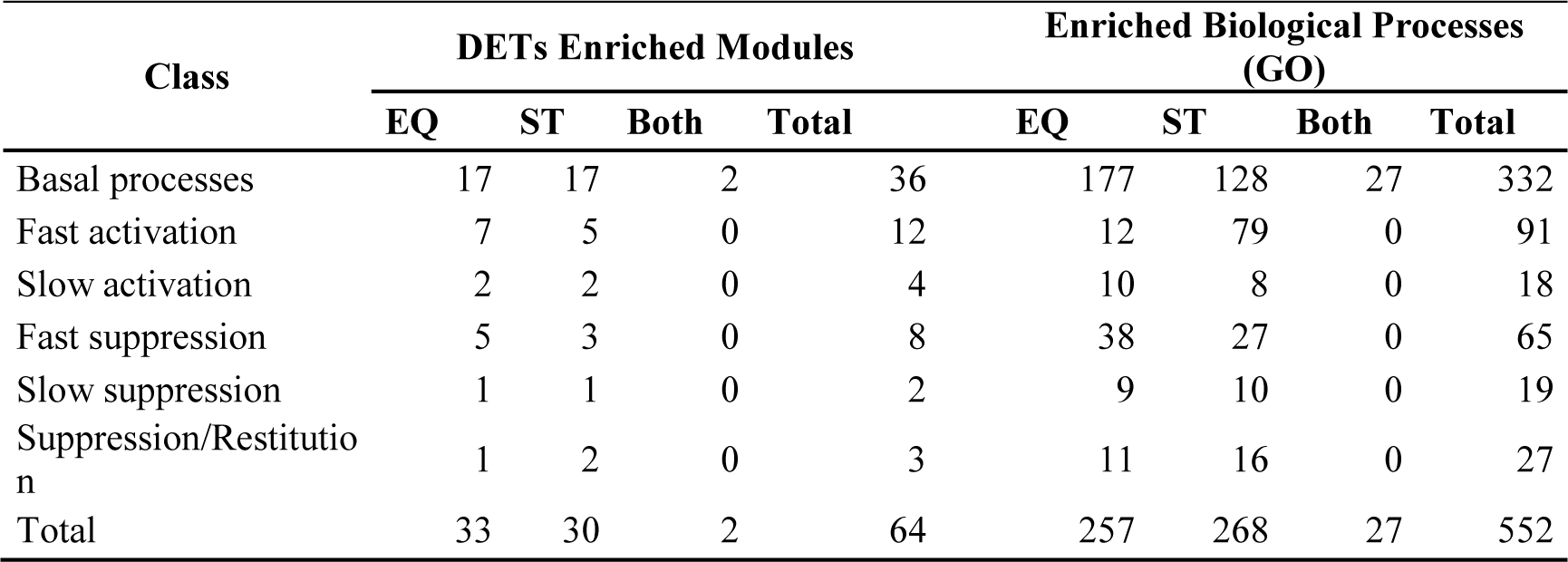
Temporal and functional behavior of modules enriched with DETs during the freezing shock in *A. schaueriana* seedlings. Number of modules per temporal class and biological processes enriched in the EQ and ST groups.

| Class | DETs Enriched Modules |  |  |  | Enriched Biological Processes (GO) |  |  |  |
| --- | --- | --- | --- | --- | --- | --- | --- | --- |
|  | EQ | ST | Both | Total | EQ | ST | Both | Total |
| Basal processes | 17 | 17 | 2 | 36 | 177 | 128 | 27 | 332 |
| Fast activation | 7 | 5 | 0 | 12 | 12 | 79 | 0 | 91 |
| Slow activation | 2 | 2 | 0 | 4 | 10 | 8 | 0 | 18 |
| Fast suppression | 5 | 3 | 0 | 8 | 38 | 27 | 0 | 65 |
| Slow suppression | 1 | 1 | 0 | 2 | 9 | 10 | 0 | 19 |
| Suppression/Restitution | 1 | 2 | 0 | 3 | 11 | 16 | 0 | 27 |
| Total | 33 | 30 | 2 | 64 | 257 | 268 | 27 | 552 |

### Functional enrichment of modules via gene ontology (GO)

In total, 552 enriched GO terms related to biological processes were identified, with the class *“basal processes”* having the highest number (332) and *“slow activation”* having the lowest (19) (Table 1). In general, the EQ group presented more enriched terms, except for *fast activation* and *suppression,* followed by *restitution*, which had more terms in the ST group. Individual interpretation of the enrichment of each module resulted in 12 modules with highlighted importance under freezing conditions. The highlighted modules were 32, 35, 69, 102, 128 and 183, which were connected to the EQ group, and 02, 29, 31, 120, 138 and 193, which were connected to the ST group.

### Basal processes

In this class, processes in common with DETs, such as biotic interaction, defense against pathogens, development of reproductive structures, ABA metabolism, AIA and vacuolar ion sequestration, were enriched (Supplementary Tables S5 to S8). There were also unique processes, such as the control of cell division and assembly of photosystem 2 (Supplementary Figures F3, F4 and F5).

### Fast activation

In the EQ group, in general, biological processes linked to growth and cell wall organization, response to abiotic and biotic stresses, hormonal signaling (ABA, AIA and SA), metabolic regulation via transcription, ubiquitination, and respiratory bursts were enriched. Specifically, module 35 showed enrichment of lipid and flavonoid synthesis in response to abiotic stress (Supplementary Figure F6).

In the ST group, in general, the enriched biological processes were linked to the formation of reproductive and vegetative structures; photoperiodism; methylation and demethylation of histones H3-K4; transmembrane transport of zinc and auxin; and protein phosphorylation and transposition via RNA. Specifically, module 2 showed enrichment of lipid synthesis associated with ion homeostasis and transcriptional regulation (Supplementary Figure F7).

### Slow activation

In the EQ group, in general, genes associated with biological processes linked to the development of reproductive structures, chitin catabolism, defense response deactivation, response to ozone and phytol metabolism were enriched. Specifically, phytol metabolism was enriched in module 128 (Supplementary Figure F8).

In the ST group, in general, the enriched processes were linked to cell wall organization, cellulose catabolism, primary root development, cell growth and AIA conjugation. AIA conjugation was specifically linked to module 192 (Supplementary Figure F9).

### Fast suppression

In the EQ group, in general, the enriched biological processes were linked to the synthesis of alkaloids and mucilage, regulation of transcription, biotic interaction and transmembrane transport of water and hydrogen peroxide. Specifically, module 69 was enriched in phospholipid catabolism, and module 32 was enriched in ETH signaling activation (Supplementary Figure F10).

In the ST group, the enriched biological processes were generally linked to lateral root formation, secondary growth, transcriptional regulation for heat, reproductive development, RNA transposition, carbohydrate metabolism and fucose assimilation. Specifically, module 29 was linked to secondary growth, and module 120 was linked to the activation of AIA signaling (Supplementary Figure F11).

### Slow suppression

In the EQ group, in general, the enriched biological processes were linked to the response to stress caused by fungi, herbivores, salinity and drought. Processes linked to terpene synthesis and glutamine secretion were also enriched. Specifically, module 183 was enriched in terpene synthesis associated with desiccation (Supplementary Figure F12).

In the ST group, in general, the enriched genes were linked to the synthesis of cutin, galacturonate and ETH; the response to oxidative stress; H2O2 catabolism; the regulation of growth and organization of the cell wall; and systemic resistance to fungi and bacteria. Specifically, module 138 was linked to ETH biosynthesis associated with the abiotic stress response (Supplementary Figure F13).

### Suppression followed by restitution

In the EQ group, only module 102 was enriched with DETs. The enriched biological processes were linked to circadian regulation, transcription of ribosomal RNA regulation, nucleolar chromosome condensation, oligosaccharide balance, gene silencing via microRNAs and regulation of pectin synthesis (Supplementary Figure F14).

In the ST group, in general, the enriched biological processes were linked to reproductive development, response to chitin, vernalization and light variation, resistance induced by jasmonic acid (JA), synthesis of terpenes and saponins and signaling through the ETH pathway. Specifically, module 31 was related to the activation of signaling by ETH (Supplementary Figure F15).

### Specific population network and hub identification

In the network constructed for the EQ group, the detected hubs were linked to modifications in protein structure (AMSH3 and ATG4), RNA cleavage (INSTS 9), energy balance (PNC1), and the transcription factor TFIIIB 60. In the network constructed for STs, the hubs were linked mainly to modifications in DNA/RNA (ZnF64, INSTS9, Symplekins and DBR1) and ribosomal production (DdRP1S2) (Table 2, Figure 4).

**Figure 4:**
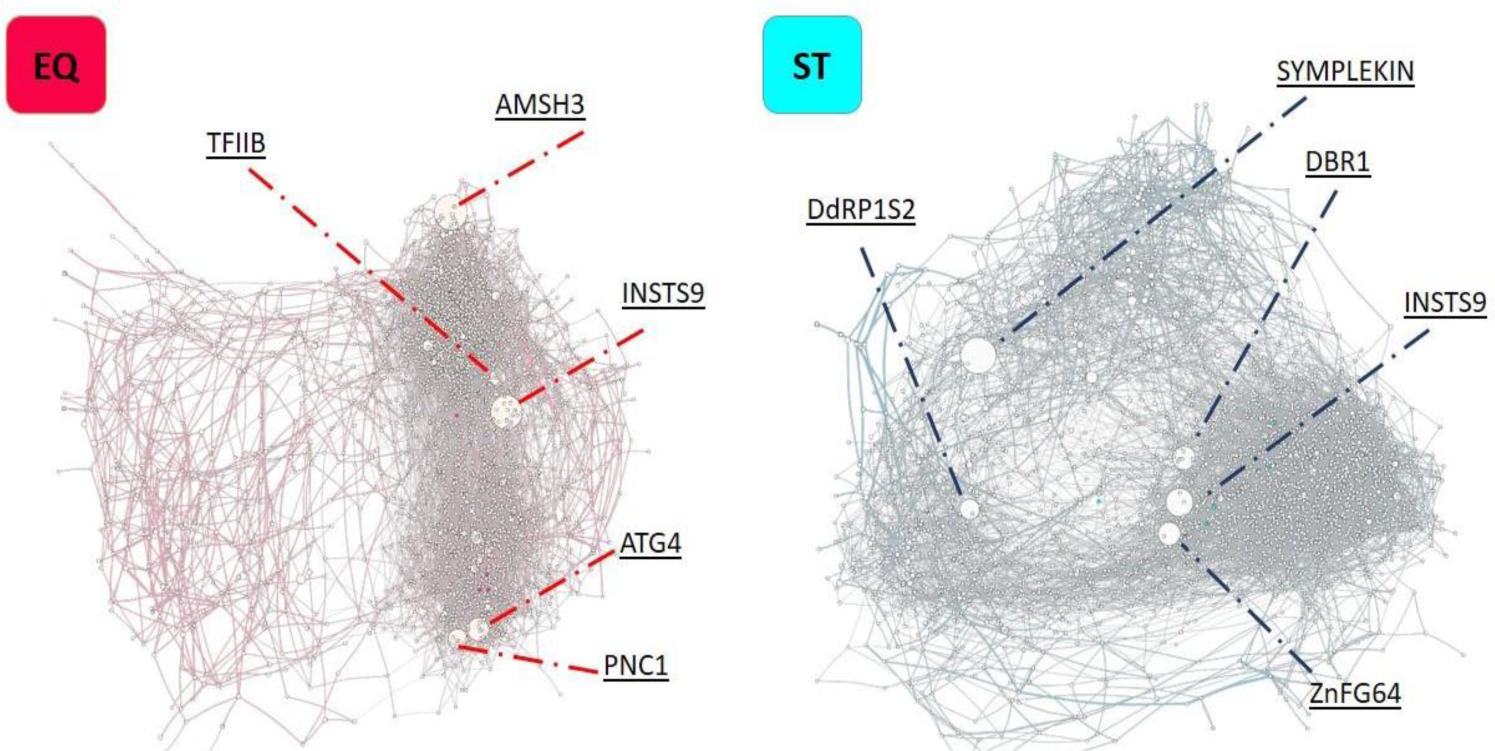
Specific networks for each EQ and ST group of *A. schaueriana* under −4°C freezing shock. Circles represent the transcripts, their size reflects the Hubscore value, and the lines represent the correlations of their expression. The hubs are highlighted and noted in each network, and the hubs are described in Table 2.

**Table 2.** Hub transcripts detected in the specific networks for each EQ and ST group under −4°C freezing of *A. schaueriana*. Functional group, Hubscore and functional transcript annotation.

| Group | Hubscore | Annotation |
| --- | --- | --- |
| EQ | 1.00 | AMSH3-like ubiquitin thioesterase 3 |
|  | 0.95 | INSTS9 - Integrator complex subunit 9 homolog |
|  | 0.79 | ATG4 -Cysteine protease ATG4 |
|  | 0.67 | PNC1- Peroxisomal adenine nucleotide carrier 1 |
|  | 0.59 | TFIIB- Transcription factor IIIB 60 kDa |
| ST | 1.00 | SYMPLEKIN |
|  | 0.90 | INSTS9 - Integrator complex subunit 9 homolog |
|  | 0.73 | ZnF64 - Zinc finger CCCH domain-containing protein 64 |
|  | 0.64 | DBR1 - Lariat debranching enzyme |
|  | 0.61 | DdRP1S2 - DNA-directed RNA polymerase I subunit 2 |

### Protein–protein interaction network

The initial network of interactions was formed by 45 proteins, with 93 and 1256 proteins identified as the 1st and 2nd neighbors, respectively, being inserted into the final network. Thus, the final network contained 1394 proteins and 1926 interactions (Figure 5). Hub proteins are linked to posttranslational modifications, such as the ubiquitination accessories SKP1, CSN5A and SUMO1; GTP-binding proteins, such as ELK4; and the membrane receptor BET12 (Table 3).

**Figure 5:**
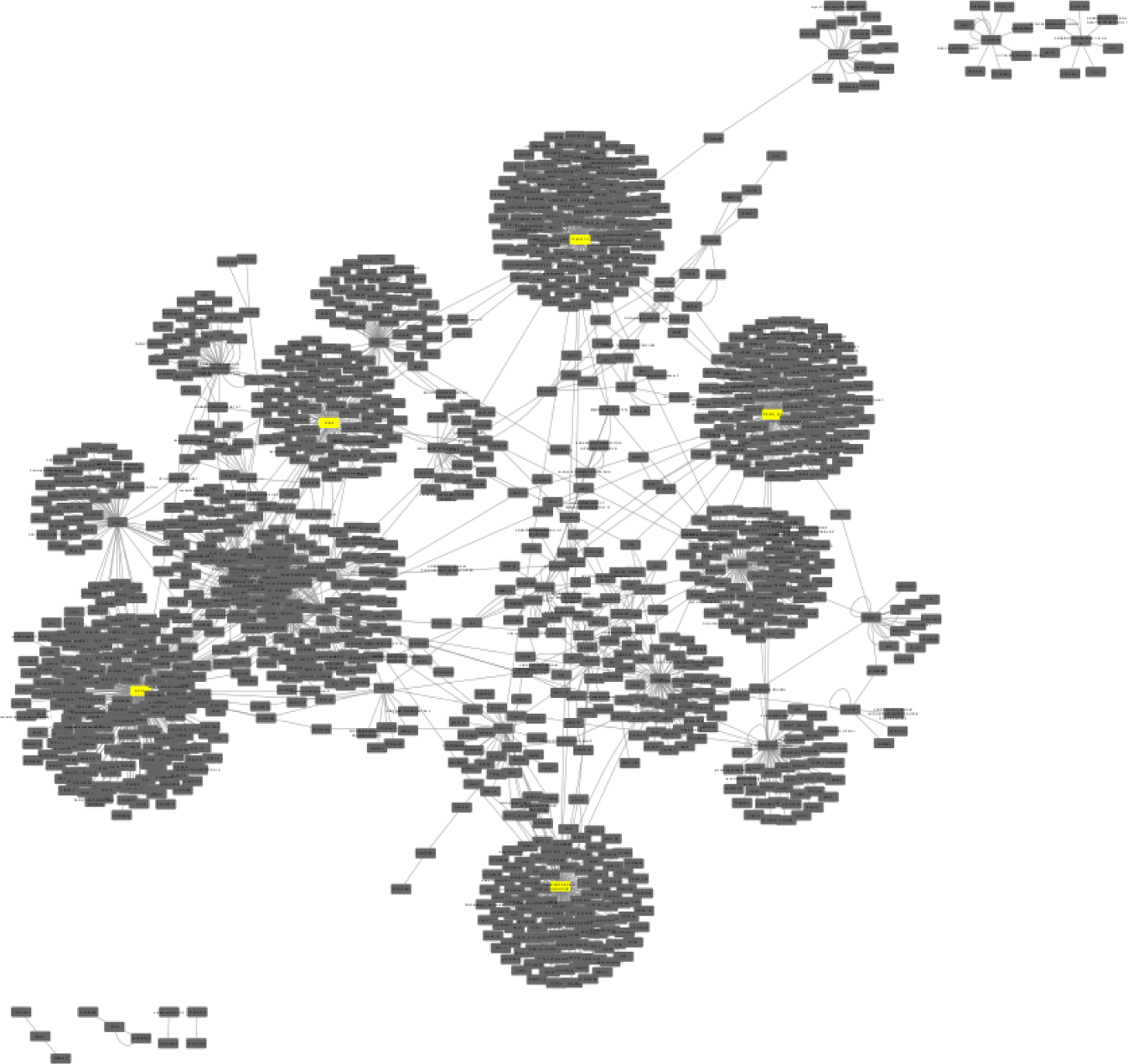
Protein–protein interaction network of *A. schaueriana* under −4°C freezing shock. The gray rectangles represent proteins, and the lines represent interactions. Hub proteins are highlighted in yellow.

**Table 3.**
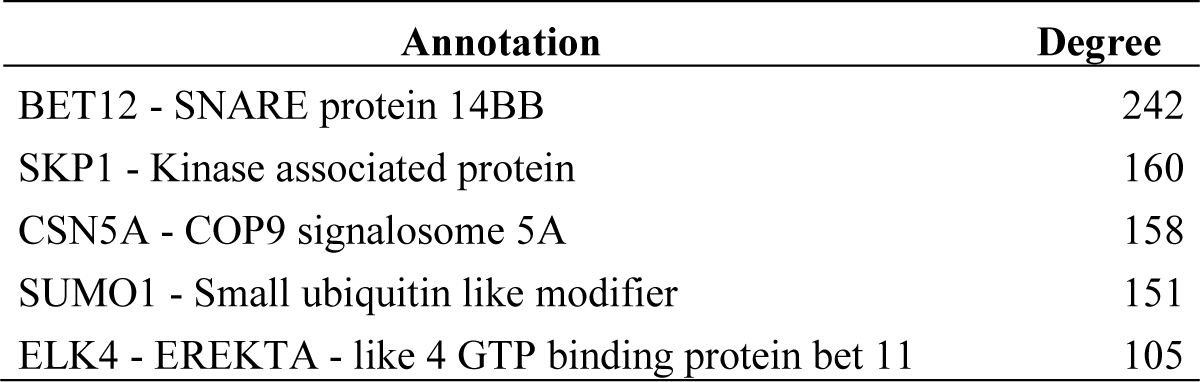
Hub proteins involved in the freezing of *A. schaueriana*. Annotation by similarity in *A. thaliana* pathways and number of connections (degree).

| Annotation | Degree |
| --- | --- |
| BET12 - SNARE protein 14BB | 242 |
| SKP1 - Kinase associated protein | 160 |
| CSN5A - COP9 signalosome 5A | 158 |
| SUMO1 - Small ubiquitin like modifier | 151 |
| ELK4 - EREKTA - like 4 GTP binding protein bet 11 | 105 |

### Differential expression validation

The nine transcripts selected for validation by RT‒qPCR were consistent with the patterns detected by differential expression analysis. The relative expression (RQ) of the genes is presented in supplementary figure F16. All the transcripts presented T test values < 0.05 when the means of the two normalizing transcripts, NR1 and NR2, were compared (Supplementary Table S13).

## DISCUSSION

In this study, we integrated a common garden experiment with differential expression approaches and coexpression networks to describe the transcriptional profile of *A. schaueriana* under freezing shock. The functional enrichment of the results allowed us to understand the direction of transcription effort during this event. Based on these results, we detected distinct patterns between plants originating from equatorial (EQ) and subtropical (ST) latitudes on the Brazilian coast. These patterns are centered on the following aspects that we will discuss below: growth and development, accumulation of cryoprotectants and antioxidants, hormonal action, posttranslational regulation, susceptibility to other stressors, lipid metabolism, and hub transcripts and proteins. Finally, we will also discuss the implications of our results for the conservation of this species and mangrove ecosystems under the current scenario of climate change.

### Temporality of results

Due to the absence of DETs in the contrasts within each EQ and ST group throughout the experiment, we used the results of the coexpression networks associated with the DETs detected in the contrasts between EQ and ST to discuss this aspect. When we checked the behavior of the samples via multidimensional scaling, we clearly observed discrete temporal groupings (Supplementary Figure F2). The low expression values probably caused an increase in the P value and low statistical reliability in these contrasts. The same does not occur when we contrast the EQ and ST samples. The basal expression profiles (time point T0) of these plants already showed marked differences, as suggested by the results of Cruz et al. 2019. Therefore, we will discuss the following topics considering that temporal changes occur in the expression patterns of plants subjected to freezing shock.

### Growth and development

After the control time point, both the EQ and ST groups showed enrichment of biological processes among the DETs linked to the development of reproductive structures, and this enrichment persisted throughout the freezing treatment (Supplementary Tables S3 to S8). This indicates that exposure to freezing temperatures did not affect this aspect of development in plants. There was also enrichment of biological processes related to the development of reproductive structures in all functional module classes in both groups (Supplementary Figures F3 to F15). However, all the plants from the EQ group produced flowers, even those from the other groups not subjected to freezing shock. However, in the ST treatment, there was no flowering of any plants. This fact indicates that the accelerated flowering in EQ may be linked to a factor that has exclusively affected plants in this group.

In the EQ group, biological processes enriched among DETs related to reproductive aspects, such as “response to light stimulus” and “post embryonic development”, were linked to transcripts annotated as LIGHT DEPENDENT short hypocotyls (LSHs). LSH is responsible for mediating reproductive development in a photoperiodic manner in *A. thaliana* (Zhao et al. 2004) and Brassica spp. (Dong et al. 2014). There were also enriched biological processes related to the development of reproductive structures and circadian regulation in the *slow suppression* and *suppression/restitution* classes (Supplementary Figures F12 and F14). These results indicate that the difference between the length of days and nights between the place of origin (Bragança-PA 0°49’S) of EQ and the greenhouse (Campinas-SP 22°54’S) may have accelerated flowering. The same trend did not occur in the ST, probably due to the smaller latitudinal shift in relation to Laguna-SC 28°S.

However, vegetative growth had opposite impacts on the two populations. The EQ population exhibited many enrichment processes linked to cell division, meristem maintenance and growth regulation among DETs (Supplementary Tables S5 and S7) after exposure to −4°C. These processes are related to genes such as PIL5-protein PIN-like5 and C78A5-cytochrome P450. The first acts by facilitating the cellular efflux of auxins, delaying plant growth, as observed in *A. thaliana* (Zazimalhová et al. 2007). The second stimulates floral development in *A. thaliana* (Ericksson et al. 2010). Enriched processes linked to growth regulation were also present in the *rapid activation* and *suppression* followed by *restitution* class (Supplementary Figures F6 and F14), another indication of the tendency to reduce vegetative growth after freezing.

On the other hand, among the DETs in the ST group, a biological process related to primary growth was identified as “stele development”, whereas not all DETs were related to growth inhibition. The “stele development” process is associated with the LONESOME HIGHWAY (LWH) gene, whose function is to regulate the formation of primary conductive tissues in *A. thaliana* (Ohashi-Ito & Bergmann 2007, Ohaish-Ito et al. 2013). Among the functional modules were enriched biological processes related to secondary growth, the formation of vegetative structures and AIA conjugation in the *fast activation, slow activation* and *suppression* followed by *restitution* (Supplementary Figures F7, F9 and F15) classes. Therefore, cold freezing had opposite effects on vegetative growth in the EQ and ST groups, suppressing and stimulating vegetative growth, respectively.

The pattern of growth inhibition associated with the stimulation of sexual reproduction detected in the EQ group may be linked to an evolutionary strategy to overcome highly unfavorable environmental conditions. In this group, several signs of stress, especially the strong action of abscisic acid (ABA), will be discussed in subsequent sections. Briefly, ABA signals plants under stress and tends to limit their growth, redirecting energy expenditure toward the production of osmolytes and saving water (Hauser et al. 2011, Kumar et al. 2017). In this situation, sexual reproduction can increase genetic variability in the next generation, thus increasing the potential for survival under adverse conditions such as rapid environmental changes (Crow 1992), which in this study corresponded to freezing shock.

### Accumulation of cryoprotectants and antioxidants

Low temperatures affect plant cells in several ways, such as by crystallizing membranes, forming ice crystals in the cytoplasm and triggering large concentrations of superoxide species (ROS) inside the cell (Heidarvand & Maali Amiri 2010). The accumulation of substances such as amino acids, peptides, carbohydrates and pigments can mitigate the effects of these impacts in a nonenzymatic way by stabilizing cell membranes, neutralizing ROS and preventing desiccation caused by freezing (Close & Beadle 2003, Heidarvand & Maali Amiri 2010, Dionne et al. 2001, Rai et al. 2002, Pan et al. 2009, Deng et al. 2019, Bilska-kos et al. 2020). These substances are called cryoprotectants.

Our results indicate that transcriptional activity is related to the accumulation of cryoprotectants in both the EQ and ST groups during freezing shock. Exclusively, in the EQ group, there were signs of accumulation of carotenoids and amino acids among DETs at T1 and T2 (Supplementary Table S14). Carotenoids such as lycopene and beta carotenes have antioxidant effects and are associated with cold stress in *Citrus* plants (Pan et al. 2009, Lado et al. 2015). When accumulated in the cell, amino acids tend to reduce the osmotic potential and neutralize the action of ROS, suggesting osmotic and antioxidant effects. For example, proline accumulates under cold conditions in species such as *A. thaliana* (Nanjo et al. 1999), *Poa annua* (Dione et al. 2001) and *Magnolia wufengensis* (Deng et al. 2019). These metabolites are closely related to ABA action in cells, as carotenoids are precursors of ABA synthesis, and proline is a result of high cellular concentrations of ABA (Othman et al. 2014, Rajasheker et al. 2019). This fact indicates that the particularities of the EQ group in the response to freezing are strongly linked to the action of ABA, which is synthesized, signaling a striking feature of this group (discussed in Hormonal action).

Commonly, both the EQ and ST groups presented biological processes and enzymatic pathways related to the accumulation of anthocyanin and small carbohydrates such as sucrose, galactose (exclusive to the ST group) and raffinose (Supplementary Tables S10, S11 and S14). Anthocyanin is a flavonoid known to accumulate in plants such as *Fagopyrum tataricum* (Li et al. 2015) and *Brassica rapa* (Ahmed et al. 2015) under low temperature and has photoprotective and antioxidant effects (Close & Beadle 2003). Small-chain carbohydrates such as galactose, sucrose and raffinose also tend to accumulate in association with cold conditions. These molecules can also act by reducing oxidative and osmotic stress in *Magnolia* (Deng et al. 2019) and *Miscanthus* (Bilska-Kos et al.). 2020) and *Ziziphus* (Zhou et al. 2020).

Therefore, our results suggest that both the EQ and ST groups accumulate osmoprotective and antioxidant substances that may confer greater tolerance to low temperatures. However, these groups differ regarding the pathways activated and metabolites generated under freezing conditions. There is no evidence in the literature about any difference in the cryoprotective potential of the accumulated metabolites. The particularities linked to the EQ group are probably related to the action of ABA in these plants.

### Hormonal action

As sessile organisms, plants have developed sophisticated ways to resist adverse climate, soil, and disease conditions. Many resistance pathways to these conditions are mediated by plant hormones such as auxins (AIA), abscisic acid (ABA), salicylic acid (SA), jasmonic acid (JA), cytokinins (CK), gibberellins (GA), strigolactones (SL), brassinosteroids (BR) and ethylene (ETH). These substances can quickly alter the metabolism and physiology of plants at the cellular and systemic levels (Eremina et al. 2016).

The action of ABA is commonly linked to abiotic stress conditions, and ABA accumulation is a highly conserved mechanism among plants under unfavorable conditions and severely impacts their metabolism (Kumas et al. 2017). A positive relationship between ABA production and concentration and freezing tolerance was verified in *A. thaliana* under cold conditions. ABA may promote the influx of calcium ions and the expression of CBF/DREB/ICE genes (Thomashow 2010). The latter are transcription factors involved in the expression of cold responsive genes (CORs), which confer metabolic responses that result in the mitigation of the negative impacts of cold (Shi-Yan et al. 2014, Eremina et al. 2016). In the EQ group, ABA was the only hormone that was enriched in processes linked to its synthesis and signaling pathways, an indication that it plays an important role in the response to freezing cold (Supplementary Table S15).

Auxins (AIA) and cytokinins (CK) have functions related to cell distension and multiplication, respectively; both have an unclear relationship with the response to cold; however, plant growth tends to be inhibited under these conditions (Eremina et al. 2016). According to our results, the AIA and CK signaling pathways are enriched among DETs at T2 exclusively in the EQ. These pathways are linked to ethylene-responsive transcription factors such as AIL5-AP2-like ethylene-responsive factors for AIA, CRF4 and RAP2.4-for CKs. These transcription factors, also known as DREB/CBF, promote cold tolerance, as documented in *Populus* spp., *Brassica napus* and *Solanum lycopersicum* (Thomashow 2010). Other hormones, such as ABA, ETH and SA (Eremina et al. 2016), also affect these same transcription factors, and identifying which and how hormones affect their expression separately is difficult. Another important characteristic is that RAP2.4 and AIL5 are also related to the flowering process (Kieber & Shaller 2018), a phenomenon in which several biological processes have been enriched in plants from both groups since T0. Flowering is also stimulated by the action of gibberellins (Bao et al. 2019), whose signaling pathway is also enriched among DETs in EQ at T1.

The accumulation of ethylene (ETH) is strongly expected to occur in plants under stressful conditions, but its role in the cold response is unclear. Moreover, ETH can confer tolerance by promoting the expression of CBF/DREB genes or even repressing them when the concentration is too high. On the other hand, ETH can directly promote the expression of COR genes and confer cold tolerance (Eremina et al. 2016). The ET pathway, which is a common pathway involved in hormone action during freezing in plants from contrasting locations, was enriched in both the EQ and ST groups.

Finally, JA, SL and BR were enriched only in STs at time point T2 (Supplementary Tables S11 and S15). JA is associated with a single pathway associated with the ethylene signaling pathway, and JA can trigger the expression of CBF/DREB genes, increasing cold tolerance (Eremina et al. 2016). According to our results, the action of JA is related to the Y3148-probable LRR serine/threonine receptor-like kinase protein-encoding gene. Protein kinase receptors are very important for signaling processes and stress perception, and the activity of these proteins can trigger cellular signaling cascades and metabolic responses (Park et al. 2014). Among DETs, SLs have genes linked to their biosynthesis, such as DAD2, and their activity under cold conditions can stimulate primary growth processes and neutralize ROS, as reported in *Brassica rapa* (Zhang et al. 2020, Bhoi et al. 2021). Under cold conditions, BRs tend to induce the expression of the CBF/COR complex and promote tolerance, as observed in *A. thaliana* (Kagale et al. 2007). The application of exogenous BRs to corn (Singh et al. 2012) and pumpkin (Jiang et al. 2013) also achieved similar results.

As expected, hormonal action was strong in both groups, but different hormones, except for ETH, acted in each group. In the EQ, we detected a greater number of hormones and a more pronounced action of ABA, while in the ST, there were fewer hormones involved in the process, and ABA showed no signs of participation.

### Posttranslational changes

Plants can sense changes in the environment and quickly alter their metabolism through changes in protein structure and activity. These changes occur mainly through ubiquitination, phosphorylation and methylation (Lyzenga & Stone 2011, Serrano et al. 2018, Zeng et al. 2019). These changes can interfere with detection and signaling pathways under stressful conditions and quickly modify the dynamics of gene expression by interacting with transcription factors and other regulatory proteins (Lyzenga & Stone 2011, Serrano et al. 2018). Among DETs, in our study, the enrichment of posttranslational changes differed between groups. In the EQs, there was an enrichment of processes related to ubiquitination and “protein ubiquitination” (Supplementary Table S7), and in the STs, there was an enrichment of processes related to dephosphorylation (Supplementary Table S8), both at time point T2. Among the groups of coexpressed transcripts, the ubiquitination process was enriched in the *Rapid Activation* class of EQs. In the ST, the phosphorylation, methylation and demethylation of H3K4 histones were also enriched in the *Rapid Activation* class (Supplementary Figure F6).

The ubiquitination process consists of the addition of a ubiquitin group to proteins, changing their conformation and activity and making them accessible for proteolysis (Stone 2019). This process has been found to have different and opposite impacts on species; for example, in *A. thaliana,* its activity is both positively (Xu & Xue 2019) and negatively related to cold tolerance (Kim & Kim 2013). In other species, such as rice, the ubiquitination process is related to increased tolerance to cold, which is detrimental to increased susceptibility to water restriction (Min et al. 2016). These findings reveal the exact relationships between these genes and the cold tolerance complex.

The process of adding or removing phosphate groups to proteins also significantly changes their activity. The addition of phosphate groups is carried out by kinases in a process called phosphorylation, and the reverse process, dephosphorylation, is carried out by phosphatases (Chen et al. 2021). According to our results, the ST population at T2 (Supplementary Table S8) had TCP-alpha-alpha-trehalose-phosphate-synthase genes linked to dephosphorylation, more precisely, the TCP genes 7, 8, 10 and 11. These genes exhibit increased expression under abiotic stresses; for example, in cassava, they promote cold tolerance by inducing sucrose accumulation (An et al. 2012). The addition or removal of methyl radicals to histone tails directly reflects access to chromatin by polymerases, impacting the expression pattern of several genes. This regulatory mechanism has already been reported in *A. thaliana* (Miura et al. 2020) and potato (Zeng et al. 2019) under cold and freezing conditions.

Posttranslational processes between DETs were enriched only in T2 for both populations, but these same processes were enriched in groups coexpressed with the *fast activation* class. This indicates that the expression patterns of these genes were correlated with T1 before they showed a considerable change in expression, as occurred in T2.

### Susceptibility to other stressors

Exposure to cold can aggravate or neutralize the action of other stresses, such as drought, pathogen infection and disease, triggering different metabolic consequences. In our study, both the EQ and ST groups presented enriched biological processes among DETs related to interactions with pathogens and mechanical injuries after freezing shock (Supplementary Tables S3 to S8), indicating that cold did not interfere with these defense mechanisms. The EQs included DETs responding to anoxia, “response to anoxia”, and a lack of water, “response to water deprivation”. Among the coexpressed genes, genes related to drought and salinity responses were enriched in the *fast activation* and *slow suppression* classes (Supplementary Figures F6 and F12).

The impacts of cold and drought induce the expression of similar metabolic pathways, the CBF/DREB gene pathway and ABA concentrations (Thomashow 2010); both of these pathways can promote the accumulation of osmolytes to avoid water loss (Rajasheker et al. al. 2019). Freezing cold also reduces the availability of liquid water in the soil, in addition to the formation of ice crystals on leaves, which can dry out tissues and cause symptoms of water restriction (Verslues et al. 2006). Excess salt in the environment also contributes to osmotic imbalance, aggravating hydric conditions and increasing ABA levels (Acosta-Matos et al. 2017, Yu et al. 2020). Low oxygenation conditions can induce the formation of ROS due to anaerobic activity, increasing pressure on cellular structures (Lukatkin 2002, Das et al. 2016). The hypoxia and anoxia response pathways are mediated by ETH and its associated transcription factors, such as the ethylene responsive factor (ERF) superfamily (Zhou et al. 2020).

Among the DETs, the STs presented biological processes related to the increase in the activity of retrotransposons; DNA modifications such as “DNA integration” and “DNA recombination” at T1; and “transposition-RNA mediated” at T2 (Supplementary Table S6 and S8). DNA modifications are associated with transposon genes such as POLR1 and POLR2, which are retrovirus-related pol protein transposon REs, and YI3B, transposon TY3. The activity of these transposable elements is a common symptom in sin plants under stress conditions and may be related to the lack of control of replication and transcription mechanisms (Lanciano & Mirouse 2018).

However, in the ST, there was enrichment of the enzymatic pathway of inositol phosphate metabolism. The accumulation of these molecules and their derivatives in plants under various stresses is common (Yang et al. 2008). This process is generally regulated by ETH-sensitive transcription factors and triggers calcium ion signaling cascades. Consequently, there is an increase in tolerance to osmotic imbalances, high salinity and cold, as reported in corn, kiwi and *A. thaliana* (Nelson et al. 1998, Munik & Verner 2010, Zhan et al. 2021). Among the coexpressed genes, genes associated with the processes of response to cold and oxidative stress were enriched. Oxidative stress, as mentioned previously, is a common symptom of cold freezing in plants, increasing damage to cellular structures (Lukatkin 2002).

### Lipid and phytol metabolism

Lipids are important constituents of membranous cellular structures; these structures are highly sensitive to drops in temperature, and damage to membranes is the first symptom caused in plants by environmental cooling (Ydav 2010). In particular, phospholipids, galactolipids, glycerolipids and sphingolipids are more abundant in membranes (Nishida & Murata 1996, Huby et al. 2019). These substances have an amphipathic quality that allows structural integrity and fluidity, guaranteeing full functionality. The accumulation and remodeling of these lipids has been observed in several species at low temperatures and are even linked to increased resistance (Sanghera et al. 2011). There are indications that the replacement and reorganization of structural lipids can mitigate cold-induced membrane damage. For example, phospholipids have been found in *A. thaliana* (Wang et al. 2006), galactolipids in rice (Zheng et al. 2016), sphingolipids in grape (Kawaguchi et al. 2000), and glycerolipids in *Dendrobium* (Zhan et al. 2021).

According to our results, enriched processes related to lipid metabolism were present in the enzymatic pathways enriched in DETs and among the groups of coexpressed transcripts. In the EQ, among the coexpressed groups, lipid synthesis was enriched and associated with respiratory bursts in the *Rapid Activation* class, specifically in module 35. These biological processes are linked to the action of PAH2-phosphatidate phosphatase 2, which can contribute to the accumulation of galactolipids in chloroplasts, as observed in *A. thaliana* (Nakamura et al. 2009). Lipid catabolism was enriched in the *rapid suppression* class (Supplementary Figure F8), and ETH signaling was enriched in module 69, associated with the NPC2 and NPC6 enzymes. These enzymes are nonspecific phospholipases that hydrolyze phospholipids in plants (Wang et al. 2012). In STs, there were enriched enzymatic pathways linked to the synthesis of sphingolipids and glycerolipids in T1 and T2 (Supplementary Tables S10 and S11). Among the coexpressed groups, lipid synthesis was enriched in the *rapid activation* class (Supplementary Figure F7), associated with ionic homeostasis and transcription regulation, specifically related to module 2 and the action of SER1-Serpin1, proteins associated with the storage of lipids in *A. thaliana* (Cai et al. 2015).

Another substance possibly linked to lipid metabolism is phytol. This molecule is a precursor to lipids synthesized in chloroplasts (Ischebeck et al. 2006, Gaude et al. 2007) and originates from chlorophyll degradation caused by stressors such as darkness and high salinity (Gutrbrod et al. 2021). Only the EQ group presented phytol metabolism enrichment in the *slow activation* class, specifically in module 128 (Supplementary Figure F8). The transcripts linked to this biological process are related to the activity of acetyltransferases, as already verified for phytol synthesis in *A. thaliana* (Lippold et al. 2012). In general, lipid synthesis was enriched in the modules or enzymatic pathways of both populations after T1. However, in EQ, the only lipid source appears to be chlorophyll degradation, depleting the photosynthetic apparatus. On the other hand, in the ST plants, more lipid types and more enzymatic pathways were also enriched after T2, and there was no evidence of chlorophyll degradation.

### Transcripts hub and regulatory functions

The transcription process can be affected in several ways, such as by interference from transcription factors and regulatory proteins and changes in the most sensitive phases of the transcript maturation process. Interference in any of these steps can result in strong metabolic consequences, stimulating or inhibiting gene transcription, translation and even protein activity (Hunt et al. 2012, Hesselberth 2013, Xu & Xue 2019, Han et al. 2021). In this section, we will discuss the actions of hubs, which probably play a regulatory role in the transcriptional profile of *A. schaueriana* under freezing conditions in the EQ and ST groups. Our individual networks for each group revealed different regulatory mechanisms involved in this metabolic adaptation among the identified hubs. In the network constructed for the EQ group, the hub transcripts were linked to the ubiquitination process, energy use, reproductive processes and mRNA processing. In the ST group, the hubs were linked mainly to mRNA processing, modifications in chromatin structure and ribosomal synthesis (Table 2).

Ubiquitination, as mentioned in the posttranslational changes section, is a process capable of affecting metabolism through the inactivation or destruction of proteins with regulatory functions. This process has high selectivity for target proteins and demands energy expenditure; additionally, this process is very sensitive to stressful situations involving the action of ABA in metabolic transformations (Xu & Xue 2019). In addition, the hub AMSH3 and ATG4-cysteine protease transcripts act together with the ubiquitination complex and reportedly act in plants tolerant of high altitudes (Sharma & Deswai 2019), drought (Lee et al. 2019) and oxidative stress (Woo et al. 2013). The other hub transcripts in the EQ, such as TFIIIB and PNC1, are linked to transcriptional regulation and the loading of ATP molecules. TFIIB is commonly associated with stimulating the development of reproductive structures (Ning et al. 2021), an event in which biological processes are enriched during all experimental time points in this group (see Growth and development). Finally, PNC1 helps modulate energy levels by balancing intracellular ADT/ATP levels, and its action has been reported in plants under numerous stresses, complicating its precise interpretation (Fonseca-Pereira et al. 2018).

The production and processing of messenger RNA (mRNA) in eukaryotes is a highly complex process. The main steps after RNA polymerization are splicing and polyadenylation. In the first step, the noncoding regions (introns) are removed from the mRNA strand. In the second stage, a tail with several adenine molecules is inserted at the strand end, protecting it from the action of nucleases and enabling its translation. Any changes in the splicing or polyadenylation step can result in aberrant peptide expression, altering protein functionality and affecting the entire cellular metabolism (Bentley 2014).

In the ST population, three of the five hub transcripts, SIMPLEKIN, DBR1 and INSTS9, are linked to mRNA processing (Table 2). Simplekins are part of the protein complex responsible for the polyadenylation of mRNAs and are associated with alternative polyadenylation events in higher plants (Hunt et al. 2012). These events are reportedly associated with the control of oxidative stress in *A. thaliana* (Xing & Li 2010) and tolerance to extreme cold and heat in *Populus* (Yan et al. 2021). DBR1 encodes a mRNA-cleaving enzyme whose main activity is to remove introns, which can generate microRNAs (miRNAs) (Hesselberth 2013, Tyagi et al. 2019).

Currently, miRNAs, such as those found in rice (Wang et al. 2014), *Citrus* (Zhang et al. 2016) and wheat (Li et al. 2019), are associated with metabolic reprogramming in plants under low temperatures. Finally, the INSTS9 transcript was identified as a hub in both populations (Table 2); this gene encodes an endonuclease capable of cleaving mRNAs and generating miRNAs (Mendoza-Figueiroa et al. 2020).

The two remaining hubs in the ST were ZnF64 and DdRP1S2. The first corresponds to a Zn2+ ion-carrying protein that folds DNA/RNA, which acts to mechanically facilitate transcription (Laity et al. 2001). This protein can be induced by hormones in combination with JA (see Hormonal action) and is associated with resistance to various abiotic stressors in *A. thaliana*; specifically, under cold conditions, this protein is linked to the triggering of COR genes (Han et al. 2021). In the second, DsRPS1S2 corresponds to subunit 2 of RNA polymerase 1, the protein responsible for transcribing ribosomal RNAs. Normally, the return of ribosomal synthesis and activity are signs of the resumption of metabolism after adaptation to stressors, as described for *A. thaliana* under excessive cold and heat conditions (Zhang et al. 2022).

### Hub proteins and regulatory functions

Proteins are molecules of fundamental importance for living organisms, and their enzymatic and structural activities can directly modulate cellular metabolism. These modulations result in effector molecules being involved in the vast majority of cellular reactions. Examples of these proteins include RNA-synthesizing proteins (Hampsey 1998) and nucleosome-forming and DNA-folding proteins that modulate chromatin access and transcription (Gerlitz et al. 2009, Han et al. 2021). Proteins can also help transport substances, recognize pathogens (Llorente et al. 1991) and destroy or inactivate other proteins (Xu & Xue 2019). Therefore, understanding protein interactions among proteins themselves and other molecular types is essential for characterizing freezing response patterns.

In our protein interaction network, we identified two main hub processes related to ubiquitination and interaction with pathogens. Ubiquitination, previously mentioned in terms of *posttranslational changes* and *hub transcript* sections, interferes with metabolism by destroying other proteins. The hub proteins SKP1, JAB1 and SUMO1 are linked to this process directly and indirectly. SKP1 is an accessory component of the ubiquitination complex and assists in recruitment for proteolysis under abiotic stress conditions, as reported in wheat and soybean (Hong et al. 2013, Chen et al. 2018). Chen et al. (2018) also reported that SKP1 is induced by hormones such as ABA, AS and JA, which were enriched in the EQ and ST groups. JAB1 is a component of the COP9 signaling site, aiding recruitment for proteolysis and reducing sensitivity to AIA (Schwechheimer et al. 2001), as occurs in EQs (see Growth and development). Finally, SUMO1 acts as a positive regulator of ubiquitination (Castro et al. 2012). During the cold response process, the activity of SUMO1 is stimulated by SA (Miura & Otha 2010), and SA can induce the accumulation of anthocyanins, as reported in apple (Jiang et al. 2022).

Biological processes associated with interactions with pathogens were enriched at all time points in both the EQ and ST groups (Supplementary Tables S3 to S8), and the hub proteins related to these processes were BET12, ELK4 and SUMO1. BET12 is a type of membrane receptor protein, and its activity is linked to the formation and fusion of membranous vesicles, an essential process for the recognition and resistance of pathogens (Jahn & Scheller 2006, Chung et al. 2018). ELK4 is a receptor kinase that acts specifically during resistance responses to pathogens (Llorente et al. 2005). SUMO1 was also reported to be involved in the SA-mediated immune response in *A. thaliana* (Ingole et al. 2021).

### Challenges to conservation in A. schaueriana

Coastal areas are the most populated by humanity, where high rates of territorial occupation, economic development and local ecosystems are severely threatened (He & Silliman 2019). Mangroves are found precisely in these areas, and estimates indicate that up to 80% of their natural areas have already been suppressed (FAO 2003). Currently, such areas are threatened by alarming rates of deforestation—approximately 2% per year (FAO 2003). The destruction of mangroves generates disastrous environmental and climatic consequences due to the high level of CO2 released during the process, the interruption of the life cycle of marine species and biogeochemical processes (Duke et al. 2007, He & Silliman 2019). In this scenario, mangrove conservation is an urgent and challenging task, and restorative actions must be effective and scientifically based, aiming for assertiveness.

The Brazilian coast plays an important role in this issue; this country contains most of the 2 million ha of South American mangroves (FAO 2007). This amount corresponds to 9% of the carbon sequestration of the world’s mangroves (Hamilton & Friess 2018). In these areas, there is knowledge about patterns of genetic and functional diversity of the main tree species in the region, such as *Rhizophora mangle*, *A. germinans* and *A. schaurianna* (Mori et al. 2015, Bajay et al. 2018, Curz et al. 2019 and 2020). Two locally adapted genetic/functional groups were identified, the first linked to the equatorial latitudes of the coast and the second to the subtropical latitudes. These groups are characterized by allelic composition, gene expression and specific morphological traits and are considered distinct evolutionary subunits. Our results reiterate this premise for the freezing response aspect, in which the subtropical group (ST) of *A. schaueriana* shows signs of decreased susceptibility, such as growth maintenance and metabolic resumption (see growth and development and transcript hubs). Therefore, we recommend that the management and conservation of these areas be carried out on an individual basis.

Mangrove restoration is a highly complex and costly process, and plant survival is closely linked to postplanting care; additionally, mangrove restoration has a greater chance of success on small area scales (Kodikara et al. 2017, Lovelock et al. 2022). Problems with the establishment of seedlings in the early stages are mainly due to soil instability, flooding, erosion and thermal fluctuations (Firess et al. 2022). This fact highlights the need for interdisciplinary knowledge and the search for genotypes suited to the specificities of each location. In this context, the selection of genotypes for eventual restoration programs must include survival in the greatest number of possible future scenarios, thus maintaining the evolutionary potential of the restored areas.

Although future climate estimates point to an increase in warming and aridity on the Brazilian coast (Vidal et al. 2021), we reiterate the need to assess the possible survival of individuals from extreme freezing events. Our results provide support for more assertive restorative actions, and we propose here that these actions be used for direct restoration programs. This is the first expression profile study of Brazilian mangroves directed to a specific stressor. Similar studies with other stressors can help to further refine management and conservation actions in mangroves on the South American Atlantic coast.

## CONCLUSIONS

This study allowed us to describe the molecular response of *A. schaueriana* to freezing shock and the divergent behavior between plants from Brazilian equatorial (EQ) and subtropical (ST) localities (Figure 6). In EQ, the response is strongly based on the action of ABA and stress signals throughout the experiment. This process negatively impacts plant growth and promotes the accumulation of carotenoids, anthocyanins and lipids through chlorophyll degradation. On the other hand, in the ST, there are fewer hormones active in the process and signs of primary growth and metabolic normalization. The accumulation of substances is mainly based on sucrose, anthocyanins and lipids, the latter not dependent on chlorophyll degradation. Based on these results, we hypothesize that susceptibility to freezing damage is greater in EQ plants than in ST plants. Therefore, we recommend that this fact be accounted for when managing this species, especially at higher latitudes, which are more prone to lower temperatures and extreme freezing events.

**Figure 6:**
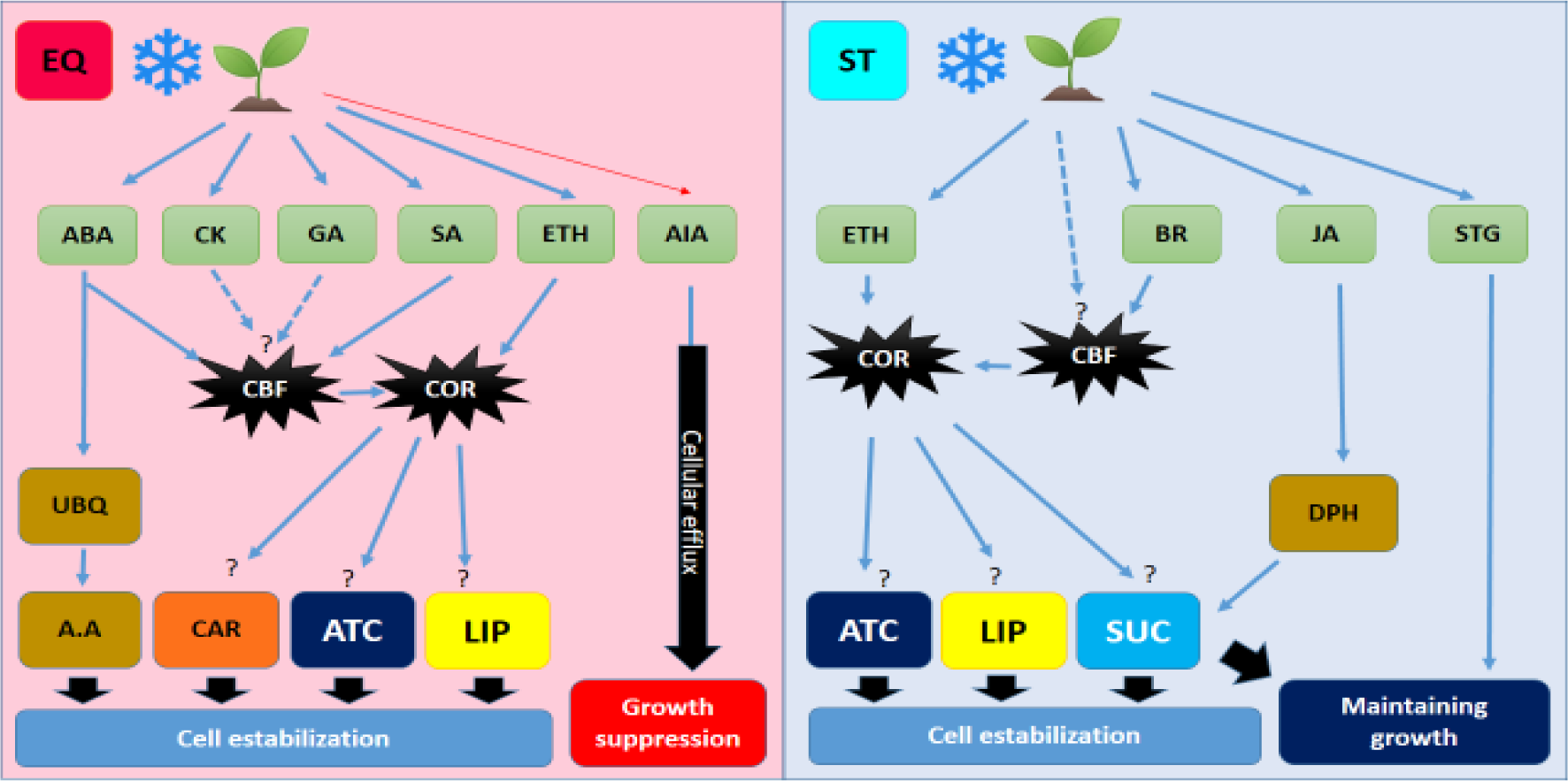
Representative scheme of the response to freezing shock (−4°C) in *A. schauriana* seedlings from equatorial (EQ) and subtropical (ST) populations on the Brazilian coast. The blue arrows indicate a positive relationship between the phenomenon or substance and possible physiological consequences. The red arrows indicate a negative relationship, and the dashed arrows indicate possible relationships according to the literature. ABA – abscisic acid, BR – brassinosteroid, SA – salicylic acid, JA – jasmonic acid, CK – cytokinin, AIA – auxin, SL – strigolactone, ETH – ethylene, UBQ – ubiquitination, DPH – dephosphorylation, ATC – anthocyanin, A.A – amino acids, CAR – carotenoids, LIP – lipids. CBF and COR – Complexes of genes and transcription factors most commonly associated with the response to cold and freezing in plants.

## ACKNOWLEDGMENTS

This work was supported by grants from the São Paulo Research Foundation (FAPESP), the Conselho Nacional de Desenvolvimento Científico e Tecnológico (CNPq), and the Coordenação de Aperfeiçoamento de Pessoal de Nível Superior (CAPES, Computational Biology Program). **YAM** received PhD fellowships from (CAPES/PROEX (88887.373880/2019–00) and FAPESP (2019/21100-0). **AAP** received a postoctoral fellowship from FAPESP (2018/00036-9). **JDV** received an Alexander von Humboldt Climate Protection Fellowship (2022-2023 cohort). **AHA** received a PhD fellowship from FAPESP (2019/03232-6). **MVC** received a PhD fellowship from FAPESP (2013/26793-7) and the CAPES Computational Biology Program (PhD SWE 8084/2015-07, PhD 88887.177158/2018–00). **APS** received a research fellowship from CNPq 312777/2018-3.

