## Supplementary tables and figures for "Molecular attributes of the tropical tree *Avicennia schaueriana* involved in the response and tolerance to low temperatures"

### SUPPLEMENTARY MATERIAL

#### *Supplementary tables*

**Table S1** - Structural aspects of the de novo assembly of the A.schauerianna transcriptome under -4ºC freezing shock.

| Parameter | Total Transcritpome | Unigene |
|---|---|---|
| N10 (pb) | 6014 | 5184 |
| N20 (pb) | 4700 | 3911 |
| N30 (pb) | 3925 | 3097 |
| N40 (pb) | 3326 | 2425 |
| N50 (pb) | 2826 | 1822 |
| Median (pb) | 1122 | 458 |
| Average (pb) | 1659 | 930 |
| GC % | 39.59 | 39.59 |
| Total bases assembled | 433 mi | 98 mi |
| Total *contigs* | 261312 | 106338 |

**Table S2 –** Completeness of the de novo assembly of the *A.schauerianna* transcriptome under low temperatures. Universal orthologs shared among the seed plant clade (embryophites) annotated via BUSCO.

| BUSCOs anotados | Groups | (%) |
|---|---|---|
| Complet | 691 | 50.3 |
| Complet single copies | 676 | 49.2 |
| Complet and duplicated | 15 | 1.1 |
| Fragmentad | 467 | 34 |
| Missing | 217 | 15.7 |
| Total de BUSCOs avaliados | 1375 | 100 |

**Table S3 –** GO terms associated with biological processes enriched in samples from the Equatorial group (EQ) of *A. schaueriana* at time T0. Term identifier (GO ID), description, number of terms noted in the transcriptome, number of significant terms in this group and time, number of expected terms and p-value obtained by Fischer's test.

| GO ID | GO term | Noted | Significant | Expected | p-value |
|---|---|---|---|---|---|
| GO:0006952 | *defense response* | 2498 | 56 | 29.99 | 5.80E-05 |
| GO:0050832 | *defense response to fungus* | 466 | 16 | 5.59 | 8.60E-05 |
| GO:0052576 | *carbohydrate storage* | 9 | 3 | 0.11 | 0.00014 |
| GO:0009836 | *fruit ripening, climacteric* | 9 | 3 | 0.11 | 0.00014 |
| GO:0006949 | *syncytium formation* | 25 | 4 | 0.3 | 0.00021 |
| GO:0055114 | *oxidation-reduction process* | 1908 | 35 | 22.9 | 0.00033 |
| GO:0009800 | *cinnamic acid biosynthetic process* | 13 | 3 | 0.16 | 0.00045 |
| GO:0010216 | *maintenance of DNA methylation* | 16 | 3 | 0.19 | 0.00085 |
| GO:0048579 | *negative regulation of long-day photoperiodism, flowering* | 16 | 3 | 0.19 | 0.00085 |
| GO:0034720 | *histone H3-K4 demethylation* | 5 | 2 | 0.06 | 0.0014 |
| GO:0010483 | *pollen tube reception* | 23 | 3 | 0.28 | 0.00254 |
| GO:0006559 | *L-phenylalanine catabolic process* | 25 | 3 | 0.3 | 0.00324 |
| GO:0006817 | *phosphate ion transport* | 126 | 6 | 1.51 | 0.00423 |
| GO:1902438 | *response to vanadate(3-)* | 9 | 2 | 0.11 | 0.00489 |
| GO:0031408 | *oxylipin biosynthetic process* | 59 | 4 | 0.71 | 0.00552 |
| GO:0055085 | *transmembrane transport* | 1775 | 28 | 21.31 | 0.00754 |
| GO:0080144 | *amino acid homeostasis* | 12 | 2 | 0.14 | 0.00876 |
| GO:0006355 | *regulation of transcription, DNA-template* | 2560 | 38 | 30.73 | 0.00896 |
| GO:0009788 | *negative regulation of abscisic acid-activity* | 107 | 5 | 1.28 | 0.0095 |

**Table S4** – GO terms associated with biological processes enriched in samples from the Subtropical group (ST) of *A. schaueriana* at time T0. Term identifier (GO ID), description, number of terms noted in the transcriptome, number of significant terms in this group and time, number of expected terms and p-value obtained by Fischer's test.

| GO ID | GO term | Noted | Significant | Expected | p-value |
|---|---|---|---|---|---|
| GO:0018171 | *peptidyl-cysteine oxidation* | 3 | 2 | 0.02 | 0.00018 |
| GO:0010411 | *xyloglucan metabolic process* | 79 | 6 | 0.62 | 0.00022 |
| GO:0006656 | *phosphatidylcholine biosynthetic process* | 73 | 5 | 0.57 | 0.00028 |
| GO:0070072 | *vacuolar proton-transporting V-type ATPase* | 5 | 2 | 0.04 | 0.0006 |
| GO:0006580 | *ethanolamine metabolic process* | 5 | 2 | 0.04 | 0.0006 |
| GO:0070483 | *detection of hypoxia* | 5 | 2 | 0.04 | 0.0006 |
| GO:0043181 | *vacuolar sequestering* | 6 | 2 | 0.05 | 0.0009 |
| GO:0032119 | *sequestering of zinc ion* | 7 | 2 | 0.05 | 0.00125 |
| GO:0042425 | *choline biosynthetic process* | 29 | 3 | 0.23 | 0.0015 |
| GO:0019563 | *glycerol catabolic process* | 8 | 2 | 0.06 | 0.00166 |
| GO:0006121 | *mitochondrial electron transport, succinate* | 13 | 2 | 0.1 | 0.00452 |
| GO:0006952 | *defense response* | 2498 | 26 | 19.6 | 0.0046 |
| GO:0055114 | *oxidation-reduction process* | 1908 | 25 | 14.97 | 0.00694 |
| GO:0009651 | *response to salt stress* | 1012 | 17 | 7.94 | 0.00748 |
| GO:0090660 | *cerebrospinal fluid circulation* | 1 | 1 | 0.01 | 0.00785 |
| GO:0019647 | *formaldehyde assimilation via ribulose m...* | 1 | 1 | 0.01 | 0.00785 |
| GO:0010080 | *regulation of floral meristem growth* | 1 | 1 | 0.01 | 0.00785 |
| GO:0010081 | *regulation of inflorescence meristem growth* | 1 | 1 | 0.01 | 0.00785 |
| GO:1900067 | *regulation of cellular response to alkal...* | 1 | 1 | 0.01 | 0.00785 |
| GO:0019748 | *secondary metabolic process* | 467 | 8 | 3.66 | 0.00844 |
| GO:0007035 | *vacuolar acidification* | 19 | 2 | 0.15 | 0.0096 |

**Table S5 –** GO terms associated with biological processes enriched in samples from the Equatorial group (EQ) of *A. schaueriana* at time T1. Term identifier (GO ID), description, number of terms noted in the transcriptome, number of significant terms in this group and time, number of expected terms and p-value obtained by Fischer's test. * processes already enriched at time T0 (control).

| GO ID | GO term | Noted | Significant | Expected | p-value |
|---|---|---|---|---|---|
| GO:0010200 | response to chitin | 234 | 17 | 2.99 | 1.00E-08 |
| GO:0055114 | oxidation-reduction process* | 1908 | 38 | 24.35 | 3.00E-05 |
| GO:0050832 | defense response to fungus* | 466 | 17 | 5.95 | 4.50E-05 |
| GO:0035264 | multicellular organism growth | 19 | 4 | 0.24 | 8.70E-05 |
| GO:0006952 | defense response* | 2498 | 58 | 31.88 | 0.00011 |
| GO:0009873 | ethylene-activated signaling pathway | 321 | 13 | 4.1 | 0.0003 |
| GO:1900994 | (-)-secologanin biosynthetic process | 11 | 3 | 0.14 | 0.00032 |
| GO:0010117 | Photoprotection | 27 | 4 | 0.34 | 0.00036 |
| GO:0009611 | response to wounding | 404 | 15 | 5.16 | 0.00041 |
| GO:0080144 | amino acid homeostasis* | 12 | 3 | 0.15 | 0.00042 |
| GO:0009414 | response to water deprivation | 782 | 22 | 9.98 | 0.00059 |
| GO:0009688 | abscisic acid biosynthetic process | 58 | 5 | 0.74 | 0.00087 |
| GO:0032973 | amino acid export across plasma membrane | 11 | 3 | 0.14 | 0.00095 |
| GO:0045892 | negative regulation of transcription, DNA template | 525 | 15 | 6.7 | 0.00115 |
| GO:0009751 | response to salicylic acid | 381 | 13 | 4.86 | 0.00188 |
| GO:0006355 | regulation of transcription, DNA-template* | 2560 | 54 | 32.67 | 0.00189 |

| | | | | | |
|---|---|---|---|---|---|
| GO:0009407 | toxin catabolic process | 23 | 3 | 0.29 | 0.00302 |
| GO:0033481 | galacturonate biosynthetic process | 7 | 2 | 0.09 | 0.00327 |
| GO:0009833 | plant-type primary cell wall biogenesis | 154 | 7 | 1.97 | 0.00373 |
| GO:0009904 | chloroplast accumulation movement | 26 | 3 | 0.33 | 0.00431 |
| GO:0048354 | mucilage biosynthetic process involved i... | 26 | 3 | 0.33 | 0.00431 |
| GO:0009738 | abscisic acid-activated signaling pathwa... | 536 | 18 | 6.84 | 0.00432 |
| GO:0080060 | integument development | 8 | 2 | 0.1 | 0.00432 |
| GO:0016121 | carotene catabolic process | 27 | 3 | 0.34 | 0.0048 |
| GO:0009607 | response to biotic stimulus | 2150 | 45 | 27.44 | 0.0055 |
| GO:1901812 | beta-carotene biosynthetic process | 9 | 2 | 0.11 | 0.00551 |
| GO:1902438 | response to vanadate(3-)* | 9 | 2 | 0.11 | 0.00551 |
| GO:1901177 | lycopene biosynthetic process | 10 | 2 | 0.13 | 0.00683 |
| GO:0009739 | response to gibberellin | 210 | 7 | 2.68 | 0.00712 |

**Table S6 –** GO terms associated with biological processes enriched in samples from the Subtropical group (ST) of *A. schaueriana* at time T1. Term identifier (GO ID), description, number of terms noted in the transcriptome, number of significant terms in this group and time, number of expected terms and p-value obtained by Fischer's test. * processes already enriched at time T0 (control).

| GO ID | GO Term | Noted | Significant | Expected | p-value |
|---|---|---|---|---|---|
| GO:0015074 | DNA integration | 544 | 17 | 4.15 | 1.10E-06 |
| GO:0005975 | carbohydrate metabolic process | 2715 | 30 | 20.71 | 4.60E-05 |
| GO:0009607 | response to biotic stimulus | 2150 | 20 | 16.4 | 8.30E-05 |
| GO:0018171 | peptidyl-cysteine oxidation* | 3 | 2 | 0.02 | 0.00017 |
| GO:0070483 | detection of hypoxia* | 5 | 2 | 0.04 | 0.00057 |
| GO:0070072 | vacuolar proton-transporting V-type ATPase* | 5 | 2 | 0.04 | 0.00057 |
| GO:0043181 | vacuolar sequestering* | 6 | 2 | 0.05 | 0.00085 |
| GO:0032119 | sequestering of zinc ion* | 7 | 2 | 0.05 | 0.00119 |
| GO:0019563 | glycerol catabolic process* | 8 | 2 | 0.06 | 0.00157 |
| GO:0010479 | stele development | 8 | 2 | 0.06 | 0.00157 |
| GO:0010143 | cutin biosynthetic process | 33 | 3 | 0.25 | 0.00202 |
| GO:0006656 | phosphatidylcholine biosynthetic process* | 73 | 4 | 0.56 | 0.00238 |
| GO:0010411 | xyloglucan metabolic process* | 79 | 4 | 0.6 | 0.00317 |
| GO:0006310 | DNA recombination | 621 | 12 | 4.74 | 0.00388 |
| GO:0010555 | response to manitol | 17 | 2 | 0.13 | 0.00731 |
| GO:1900067 | regulation of cellular response to alkaloid* | 1 | 1 | 0.01 | 0.00763 |
| GO:0010080 | regulation of floral meristem growth* | 1 | 1 | 0.01 | 0.00763 |
| GO:0010081 | regulation of inflorescence meristem growth*. | 1 | 1 | 0.01 | 0.00763 |
| GO:0006952 | defense response* | 2498 | 28 | 19.06 | 0.00784 |
| GO:0010229 | inflorescence development | 63 | 5 | 0.48 | 0.00796 |
| GO:0019748 | secondary metabolic process* | 467 | 7 | 3.56 | 0.00806 |
| GO:0007035 | vacuolar acidification* | 19 | 2 | 0.14 | 0.00909 |

**Table S7 –** GO terms associated with biological processes enriched in samples from the Equatorial group (EQ) of *A. schaueriana* at time T2. Term identifier (GO ID), description, number of terms noted in the transcriptome, number of significant terms in this group and time, number of expected terms and p-value obtained by Fischer's test. * processes already enriched at time T0 (control), and ** processes already enriched at T1.

| GO ID | GO Term | Noted | Significant | Expected | p-value |
|---|---|---|---|---|---|
| GO:0040009 | regulation of growth rate | 15 | 4 | 0.21 | 4.70E-05 |
| GO:0030574 | collagen catabolic process | 18 | 4 | 0.25 | 0.0001 |

| | | | | | |
|---|---|---|---|---|---|
| GO:0060774 | auxin mediated signaling pathway involve... | 2 | 2 | 0.03 | 0.0002 |
| GO:0090698 | post-embryonic plant morphogenesis | 344 | 17 | 4.86 | 0.00022 |
| GO:0009407 | toxin catabolic process** | 23 | 4 | 0.33 | 0.00028 |
| GO:0055114 | oxidation-reduction process* | 1908 | 41 | 26.97 | 0.00028 |
| GO:0046622 | positive regulation of organ growth | 10 | 3 | 0.14 | 0.00031 |
| GO:0009299 | mRNA transcription | 24 | 4 | 0.34 | 0.00033 |
| GO:0006662 | glycerol ether metabolic process | 24 | 4 | 0.34 | 0.00033 |
| GO:0048354 | mucilage biosynthetic process involved i...** | 26 | 4 | 0.37 | 0.00046 |
| GO:0060772 | leaf phyllotactic patterning | 3 | 2 | 0.04 | 0.00059 |
| GO:0009094 | L-phenylalanine biosynthetic process | 13 | 3 | 0.18 | 0.00072 |
| GO:0010088 | phloem development | 31 | 4 | 0.44 | 0.00092 |
| GO:0009416 | response to light stimulus | 1329 | 28 | 18.79 | 0.00096 |
| GO:0006355 | regulation of transcription, DNA-template* | 2560 | 52 | 36.19 | 0.00099 |
| GO:0034059 | response to anoxia | 8 | 3 | 0.11 | 0.00117 |
| GO:0045595 | regulation of cell differentiation | 228 | 8 | 3.22 | 0.00178 |
| GO:0040019 | positive regulation of embryonic develop... | 5 | 2 | 0.07 | 0.00194 |
| GO:0046256 | 2,4,6-trinitrotoluene catabolic process | 5 | 2 | 0.07 | 0.00194 |
| GO:0009873 | ethylene-activated signaling pathway** | 321 | 12 | 4.54 | 0.00221 |
| GO:1900366 | negative regulation of defense response | 6 | 2 | 0.08 | 0.00288 |
| GO:0009738 | abscisic acid-activated signaling pathway** | 536 | 19 | 7.58 | 0.00302 |
| GO:0016567 | protein ubiquitination | 951 | 25 | 13.44 | 0.00308 |
| GO:0009611 | response to wounding** | 404 | 14 | 5.71 | 0.00314 |
| GO:0016135 | saponin biosynthetic process | 22 | 3 | 0.31 | 0.00354 |
| GO:0010311 | lateral root formation | 104 | 6 | 1.47 | 0.00365 |
| GO:0009682 | induced systemic resistance | 46 | 5 | 0.65 | 0.00398 |
| GO:0080060 | integument development** | 8 | 2 | 0.11 | 0.00528 |
| GO:0006749 | glutathione metabolic process | 50 | 4 | 0.71 | 0.00543 |
| GO:1902438 | response to vanadate(3-)* | 9 | 2 | 0.13 | 0.00672 |
| GO:0010492 | maintenance of shoot apical meristem identity | 54 | 4 | 0.76 | 0.00713 |
| GO:0009944 | polarity specification of adaxial/abaxia... | 54 | 4 | 0.76 | 0.00713 |
| GO:0009736 | cytokinin-activated signaling pathway | 108 | 6 | 1.53 | 0.0086 |
| GO:0006817 | phosphate ion transport* | 126 | 6 | 1.78 | 0.00863 |
| GO:0031408 | oxylipin biosynthetic process* | 59 | 4 | 0.83 | 0.00972 |

**Table S8 –** GO terms associated with biological processes enriched in samples from the group (ST) of *A. schaueriana* at time T2. Term identifier (GO ID), description, number of terms noted in the transcriptome, number of significant terms in this group and time, number of expected terms and p-value obtained by Fischer's test. * processes already enriched at time T0 (control), and ** processes already enriched at T1.

| GO ID | GO Term | Noted | Significant | Expected | p-value |
|---|---|---|---|---|---|
| GO:0005975 | carbohydrate metabolic process** | 2715 | 60 | 28.76 | 5.40E-15 |
| GO:0042546 | cell wall biogenesis | 543 | 10 | 5.75 | 2.80E-07 |
| GO:0010411 | xyloglucan metabolic process* | 79 | 9 | 0.84 | 7.50E-07 |
| GO:0010479 | stele development | 8 | 3 | 0.08 | 6.30E-05 |
| GO:0016311 | dephosphorylation | 246 | 8 | 2.61 | 0.00026 |
| GO:0070072 | vacuolar proton-transporting V-type ATPase* | 5 | 2 | 0.05 | 0.0011 |
| GO:0043181 | vacuolar sequestering* | 6 | 2 | 0.06 | 0.00163 |

| | | | | | |
|---|---|---|---|---|---|
| GO:1901601 | strigolactone biosynthetic process | 7 | 2 | 0.07 | 0.00227 |
| GO:0032119 | sequestering of zinc ion* | 7 | 2 | 0.07 | 0.00227 |
| GO:0010229 | inflorescence development** | 63 | 6 | 0.67 | 0.00251 |
| GO:0031540 | regulation of anthocyanin biosynthetic process | 27 | 3 | 0.29 | 0.00285 |
| GO:0019563 | glycerol catabolic process* | 8 | 2 | 0.08 | 0.003 |
| GO:0009861 | jasmonic acid and ethylene-dependent sys... | 31 | 3 | 0.33 | 0.00425 |
| GO:0010078 | maintenance of root meristem identity | 32 | 3 | 0.34 | 0.00465 |
| GO:0019605 | butyrate metabolic process | 10 | 2 | 0.11 | 0.00476 |
| GO:0032197 | transposition, RNA-mediated | 66 | 4 | 0.7 | 0.00531 |
| GO:0015074 | DNA integration* | 544 | 13 | 5.76 | 0.00566 |
| GO:0071555 | cell wall organization | 1005 | 16 | 10.65 | 0.00735 |
| GO:0035434 | copper ion transmembrane transport | 13 | 2 | 0.14 | 0.00808 |

**Table S9-** Enzymatic pathways enriched in differentially expressed transcripts between the EQ and ST groups of *A. schaueriana* contrasted at the control time T0, before freezing shock at -4°C. Group, enriched pathway noted on Kegg and p value estimated via Fischer test.

| Group | Enriched enzymatic pathway | P-value |
|---|---|---|
| EQ | Biosynthesis of various plant secondary metabolites | 0.000735021 |
| | Sesquiterpenoid and triterpenoid biosynthesis | 0.003218863 |
| | Lysine degradation | 0.006349083 |
| | Carotenoid biosynthesis | 0.006349083 |
| | Plant-pathogen interaction | 0.011661473 |
| | Circadian rhythm – plant | 0.024754232 |
| | Glutathione metabolismo | 0.035665272 |
| | Zeatin biosynthesis | 0.040651542 |
| | Stilbenoid, diarylheptanoid and gingerol biosynthesis | 0.041045802 |
| | Inositol phosphate metabolismo | 0.042775967 |
| | Monoterpenoid biosynthesis | 0.048727006 |
| ST | Fatty acid elongation | 1.91618E-05 |
| | Glycerophospholipid metabolismo | 0.005287934 |
| | Oxidative phosphorylation | 0.011147102 |
| | Phagosome | 0.016103594 |
| | Plant hormone signal transduction | 0.042392261 |

**Table S10** - Enzymatic pathways enriched in differentially expressed transcripts between the EQ and ST groups *of A. schaueriana* contrasted at time T1, 15 minutes after freezing shock at -4°C. Group, enriched pathway noted on Kegg and p value estimated via Fischer test.

| Group | Enriched enzymatic pathway | P-value |
|---|---|---|
| EQ | Carotenoid biosynthesis | 0.00035789 |
| | Phenylpropanoid biosynthesis | 0.002690044 |
| | Flavonoid biosynthesis | 0.00280274 |
| | Stilbenoid, diarylheptanoid and gingerol biosynthesis | 0.007334317 |
| | Biosynthesis of various plant secondary metabolites | 0.012811382 |
| | Biosynthesis of secondary metabolites | 0.016947179 |
| | Glutathione metabolism | 0.02001334 |
| | Ribosome | 0.024018466 |
| | Tyrosine metabolism | 0.047879817 |

| | | |
|---|---|---|
| ST | Sphingolipid metabolism | 0.000639683 |
| | Galactose metabolism | 0.000703456 |
| | Glycosphingolipid biosynthesis - globo and isoglobo series | 0.000961223 |
| | Phagosome | 0.001386059 |
| | Plant hormone signal transduction | 0.001643731 |
| | Glycerolipid metabolism | 0.004140695 |
| | ABC transportes | 0.010618487 |
| | Glycerophospholipid metabolismo | 0.014700674 |
| | Oxidative phosphorylation | 0.027490599 |
| | Fatty acid elongation | 0.033339894 |
| | Glycosphingolipid biosynthesis - ganglio series | 0.039144082 |

**Table S11-** Pathways enriched in differentially expressed transcripts between the EQ and ST groups of *A. schaueriana* contrasted at time T2, 180 minutes after freezing shock at -4°C. Group, enriched pathway noted on Kegg and p value estimated via Fischer test.

| Group | Enriched enzymatic pathway | P-value |
|---|---|---|
| EQ | Glutathione metabolism | 0.00034312 |
| | Carotenoid biosynthesis | 0.001034264 |
| | Phosphatidylinositol signaling system | 0.002981948 |
| | Inositol phosphate metabolism | 0.004195933 |
| | Stilbenoid, diarylheptanoid and gingerol biosynthesis | 0.004421853 |
| | Biosynthesis of various plant secondary metabolites | 0.006863244 |
| | Flavonoid biosynthesis | 0.013293424 |
| | Phenylpropanoid biosynthesis | 0.020222175 |
| | Anthocyanin biosynthesis | 0.023239629 |
| | Diterpenoid biosynthesis | 0.031211496 |
| | Sesquiterpenoid and triterpenoid biosynthesis | 0.040094188 |
| | Biosynthesis of secondary metabolites | 0.050614243 |
| ST | Glycosphingolipid biosynthesis - globo and isoglobo series | 1.30133E-09 |
| | Galactose metabolism | 1.98253E-09 |
| | Sphingolipid metabolism | 2.8245E-08 |
| | Glycerolipid metabolism | 2.32559E-06 |
| | Fatty acid elongation | 0.001546347 |
| | Phagosome | 0.003324697 |
| | Brassinosteroid biosynthesis | 0.007994324 |
| | Inositol phosphate metabolism | 0.008227435 |
| | ABC transporters | 0.008431365 |
| | Butanoate metabolism | 0.027660933 |
| | Plant hormone signal transduction | 0.030419141 |
| | Starch and sucrose metabolismo | 0.03994159 |

**Table S12 –** Transcripts selected for validation of differential expression of *A.schaueriana* seedlings under -4°C freezing shock. Transcript code, identifier in the transcriptome, functional annotation, fold change and contrast in which it was detected as DET.

| COD | Transcript_ID | Annotatiom | LFC | CTR |
|---|---|---|---|---|
| DT2 | TRINITY_DN32157_c1_g1_i1 | PQP93298.1 - Putative disease resistance protein | (+)8.6 | T0 |
| DT3 | TRINITY_DN82976_c0_g1_i1 | XP_011076877.1 -Nuclear pore complex protein NUP1 | (+)8.15 | T0 |
| DT7 | TRINITY_DN44134_c0_g1_i3 | CCR1_ARATH - Cinnamoyl-CoA reductase 1 | (-)10 | T0 |

| | | | | |
|---|---|---|---|---|
| DT9 | TRINITY_DN41250_c0_g1_i4 | SCP40_ARATH -Serine carboxypeptidase-like 40 | (-)9 | T0 |
| DT10 | TRINITY_DN34959_c0_g1_i1 | VATB2_GOSHI - Vacuolar proton pump subunit B 2 | (-)8.12 | T0 |
| DT12 | TRINITY_DN40804_c1_g2_i10 | PIN10365.1 - Replication factor C, subunit RFC3 | (+)813 | T1 |
| DT23 | TRINITY_DN42180_c2_g1_i4 | RBL20_ARATH - Rhomboid-like protein 20 | (+)5.8 | T2 |
| DT25 | TRINITY_DN44192_c0_g1_i1 | AB11G - ABC2 transporter | (-)7.8 | T2 |
| DT26 | TRINITY_DN32601_c0_g2_i1 | GER52371.1 - pollen Ole e 1 allergen/extensin family | (-)7.7 | T2 |

**Table S13 –** Transcripts selected to validate the differential expression of *A.schaueriana* seedlings under -4°C freezing shock. Transcript code, identifier in the transcriptome, left primer, right primers and amplicon size in bases.

| COD | Transcript_ID | Primer F | Primer R | Size |
|---|---|---|---|---|
| DT2 | TRINITY_DN32157_c1_g1_i1 | CAACTGGTAACGTTGCTGCA | AGATGCGAAGAAGCTCCGAG | 80 |
| DT3 | TRINITY_DN82976_c0_g1_i1 | TGGCAAAAACGGGCTTGATG | ACCCACATCCTCTGCTCCTA | 82 |
| DT7 | TRINITY_DN44134_c0_g1_i3 | TTTGCCCTCTCAGTCGACAC | AGGCTAACACAAAGACGGCT | 111 |
| DT9 | TRINITY_DN41250_c0_g1_i4 | CATCCTGAGGCGCCATGTAA | TCCGCTCGGGGAACTACTAA | 107 |
| DT10 | TRINITY_DN34959_c0_g1_i1 | GGTGTGGGCTCCGATATACG | GCTTCGGAGGGGCTAATCAA | 93 |
| DT12 | TRINITY_DN40804_c1_g2_i10 | GTTTAGCGTGGGTTGATGGC | AGATTTCGTCCAGTTGCCGT | 84 |
| DT23 | TRINITY_DN42180_c2_g1_i4 | AACAAGGGCGCATTACTGC | CTAAGCGCATTGAGCAGGC | 88 |
| DT25 | TRINITY_DN44192_c0_g1_i1 | TGGCCCAACAGTTCACACA | TGGAGATCCAGCCTTCACCA | 107 |
| DT26 | TRINITY_DN32601_c0_g2_i1 | ATGGCTGAGGTGAGAACGC | ACTTCTGGGATGCCAAGCC | 93 |
| NR1 | TRINITY_DN44176_c0_g1_i6 | GCTCCAGGGTTAGGGTTTGT | TGACATGGCAGCCTCAAGTT | 87 |
| NR2 | TRINITY_DN42576_c0_g1_i4 | AACAGCAGCCAACCAACTTG | AGCCACAGTGCATCTGAACT | 119 |

**Table S14 –** GO terms associated with enriched biological processes linked to the production of cryoprotective substances in *A.schaueriana* plants of functional groups EQ and ST under freezing (-4°C). Cryoprotective substance, description of the term GO, functional group and time under freezing.

| Cryoprotectant | | GO Term | Group | Time |
|---|---|---|---|---|
| Amino acids | Phenylalanine | *L-phenylalaline biosinthetic process* | EQ | T2 |
| | Proline | *collagen catabolic process* | EQ | T2 |
| Carotenoids | Beta carotene | *betacatoteinoid biosybthetic process* | EQ | T1 |
| | Lycopene | *lycopene biosynthetic process* | EQ | T1 |
| Flavonoids | Anthocyanin | *regulation of antocyanin biosynthetic process* | ST | T2 |
| Carbohydrates | Sucrose | *carbohydrate metabolism* | ST | T1 |
| | | *response to manitol* | ST | T1 |

**Table S15 –** GO terms associated with enriched biological processes linked to hormonal action in *A.schaueriana* plants from functional groups EQ and ST under freezing (-4°C). Hormone, description of the GO term, functional group, functional group and time under freezing.

| Hormone | GO Term | Group | Time |
|---|---|---|---|
| ABA | *abscisic acid biosynthetic process* | EQ | T1 e T2 |
| ABA | *abscisic acid-activated signaling pathway* | EQ | T1 e T2 |

| | | | |
|---|---|---|---|
| ETH | *ethylene-activated signaling pathway* | EQ | T1 e T2 |
| SA | *response to salicylic acid* | EQ | T1 |
| GA | *response to gibberellin* | EQ | T1 |
| AIA | *auxin mediated signaling pathway involved in phyllotactic patterning* | EQ | T2 |
| CK | *cytokinin-activated signaling pathway* | EQ | T2 |
| JA | *jasmonic acid and ethylene-dependent systemic resistance* | ST | T2 |
| SL | *strigolactone biosynthesis* | ST | T2 |

*Supplementary Figures*

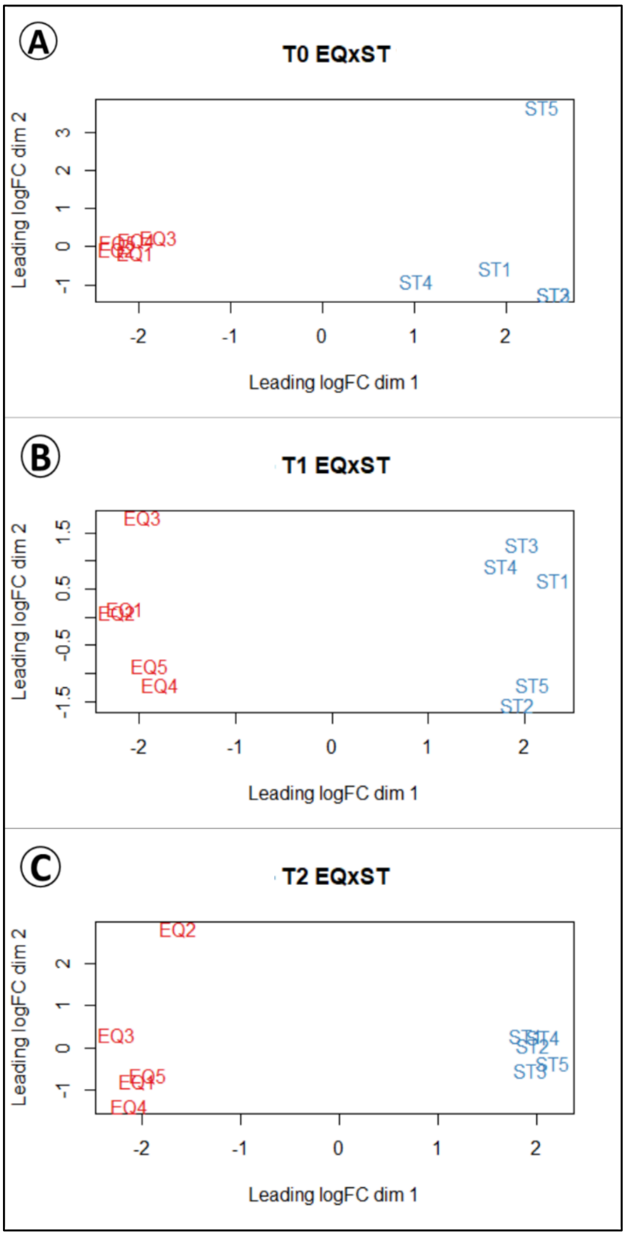


**Figure F1**- Multidimensional scaling of samples from the equatorial (EQ) and subtropical (ST) groups of A.schaueriana under freezing. Plots based on normalized expression data converted to log base 2 (LFC). Contrasts between groups within each of the times a) T0 (control), b) T1 (15 min) and c) T2 (180 min).

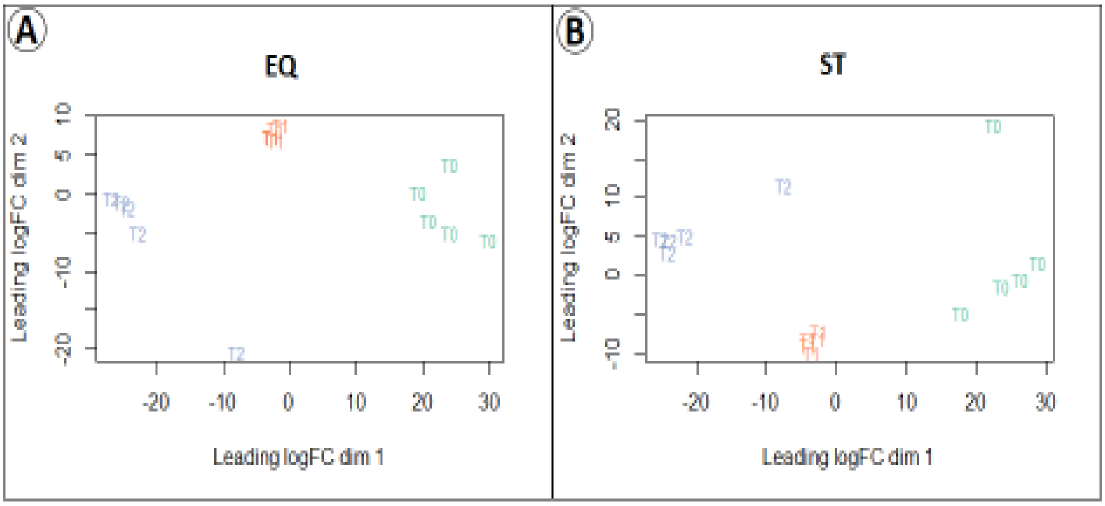


**Figure F2** – Multidimensional scaling of samples from the equatorial (EQ) and subtropical (ST) groups of *A.schaueriana* under freezing. Plots based on normalized expression data converted to log base 2 (LFC). Temporal contrasts within each of the functional groups a) EQ and b) ST.T0 (control) in green, T1 (15 min) in red and T2 (180 min) in blue.

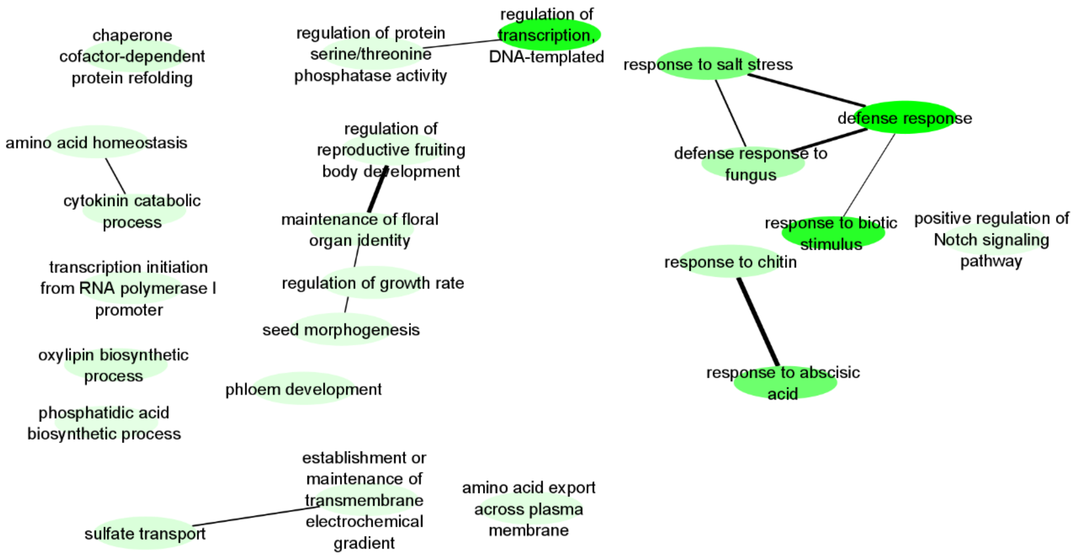


**Supplementary Figure F3 –** Summary of enriched biological processes common to the EQ and ST groups in Basal processes class of *Avicennia schaueriana*. The ellipses represent the enriched GO terms and the lines the genetic ontology relationship, the thicker the line the greater the ontogenetic relationship of the terms.

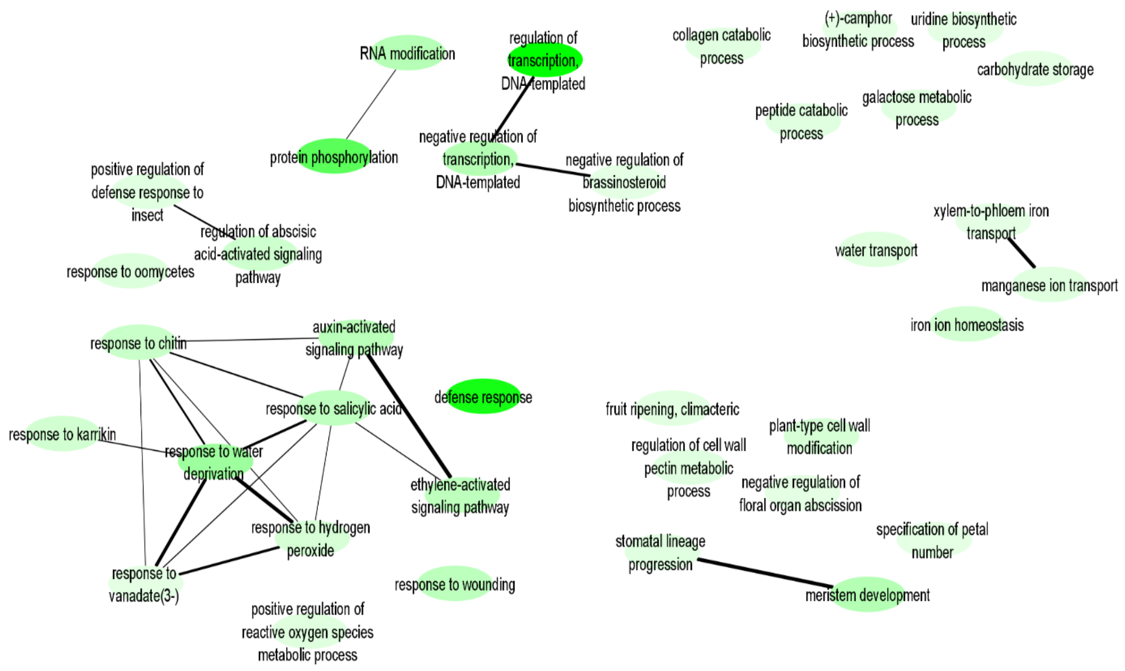


**Figure F4 –** Summary of biological processes enriched in the EQ group in Basal processes class of *Avicennia schaueriana*. The ellipses represent the enriched GO terms and the lines the genetic ontology relationship, the thicker the line the greater the ontogenetic relationship of the terms.

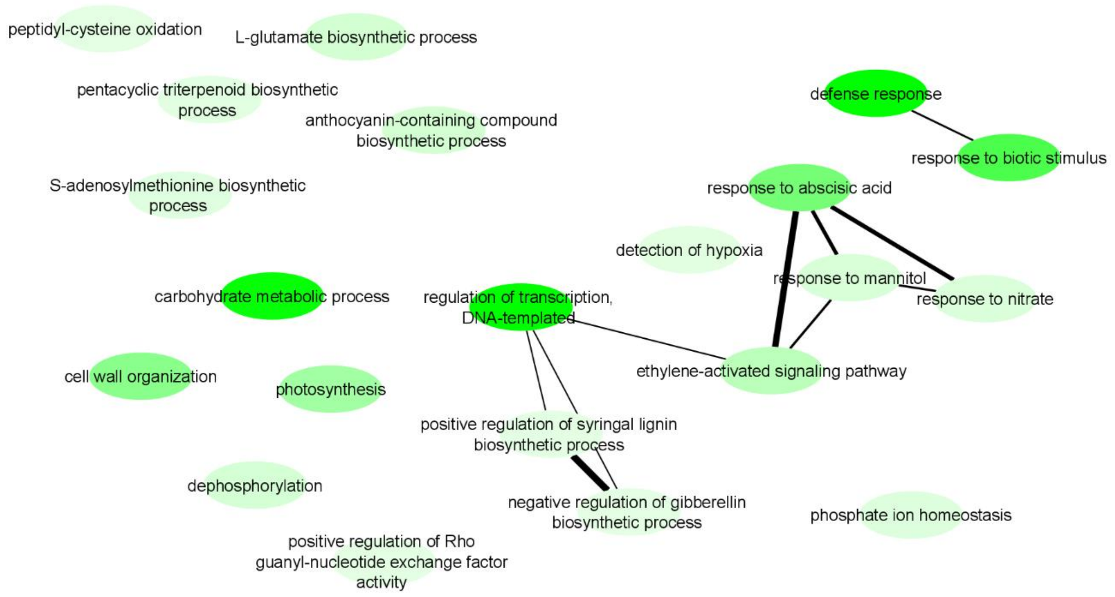


**Figure F5 –** Summary of biological processes enriched in the ST group in Basal processes class of *Avicennia schaueriana*. The ellipses represent the enriched GO terms and the lines the genetic ontology relationship, the thicker the line the greater the ontogenetic relationship of the terms. The ellipses represent the enriched GO terms and the lines the genetic ontology relationship, the thicker the line the greater the ontogenetic relationship of the terms.

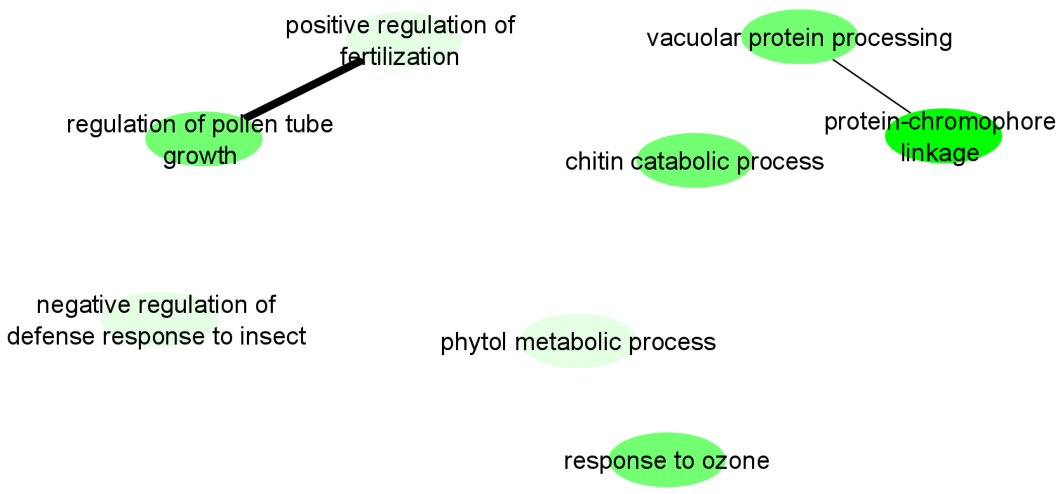


**Figure F6** – Summary of biological processes enriched in the EQ group in Rapid activation class of *Avicennia schaueriana* under freezing temperatures (-4°C). The ellipses represent the enriched GO terms and the lines the genetic ontology relationship, the thicker the line the greater the ontogenetic relationship of the terms.

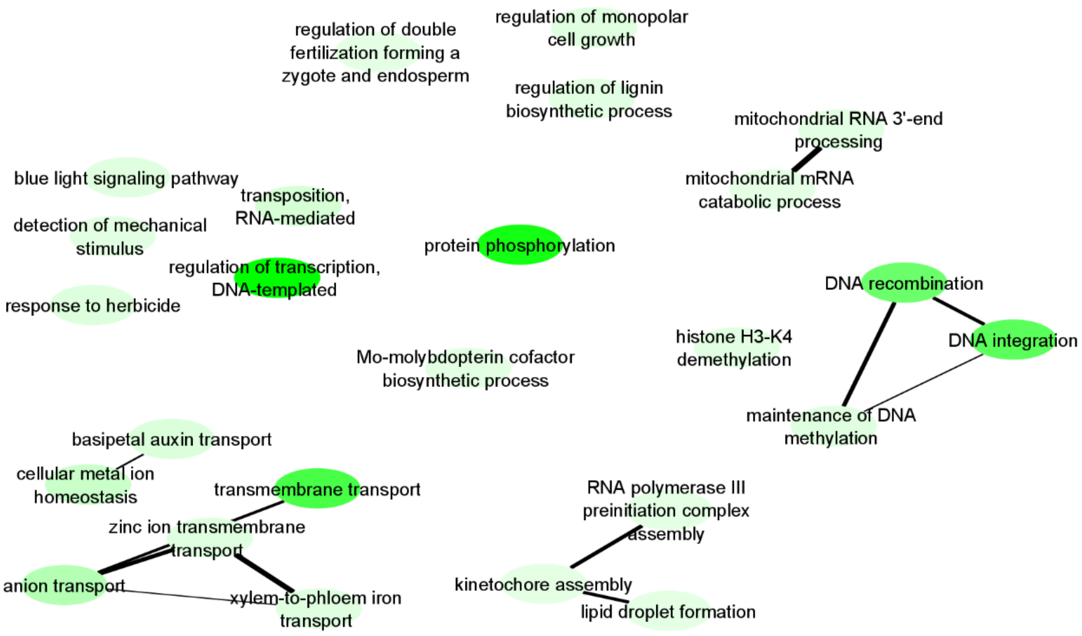


**Figure F7** – Summary of biological processes enriched in the ST group in Rapid activation class in *Avicennia schaueriana* under freezing temperatures (-4°C). The ellipses represent the enriched GO terms and the lines the genetic ontology relationship, the thicker the line the greater the ontogenetic relationship of the terms.

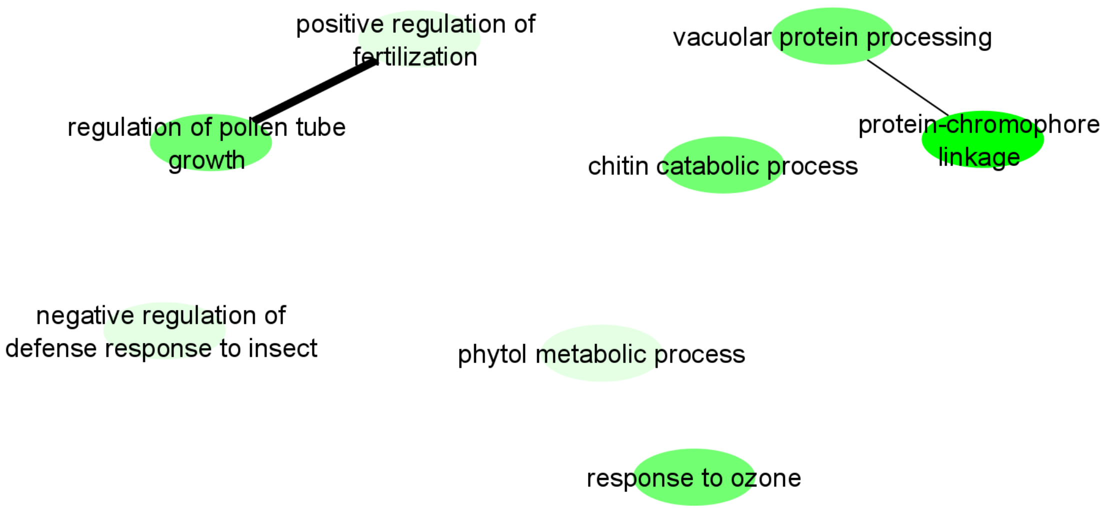


**Figure F8 –** Summary of biological processes enriched in the EQ group Slow activation class in *Avicennia schaueriana* under freezing temperatures (-4°C). The ellipses represent the enriched GO terms and the lines the genetic ontology relationship, the thicker the line the greater the ontogenetic relationship of the terms.

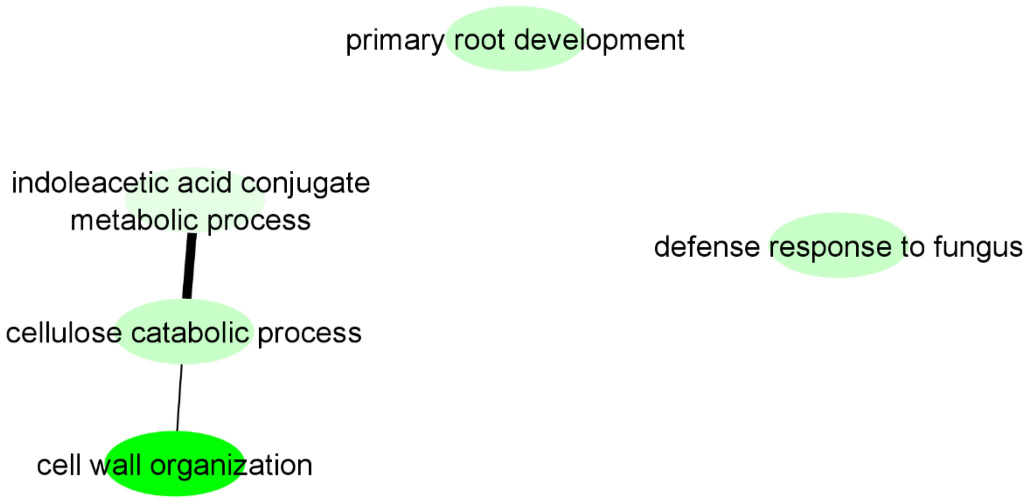


**Figure F9 –** Summary of biological processes enriched in the ST group Slow activation class in *Avicennia schaueriana* under freezing temperatures (-4°C). The ellipses represent the enriched GO terms and the lines the genetic ontology relationship, the thicker the line the greater the ontogenetic relationship of the terms.

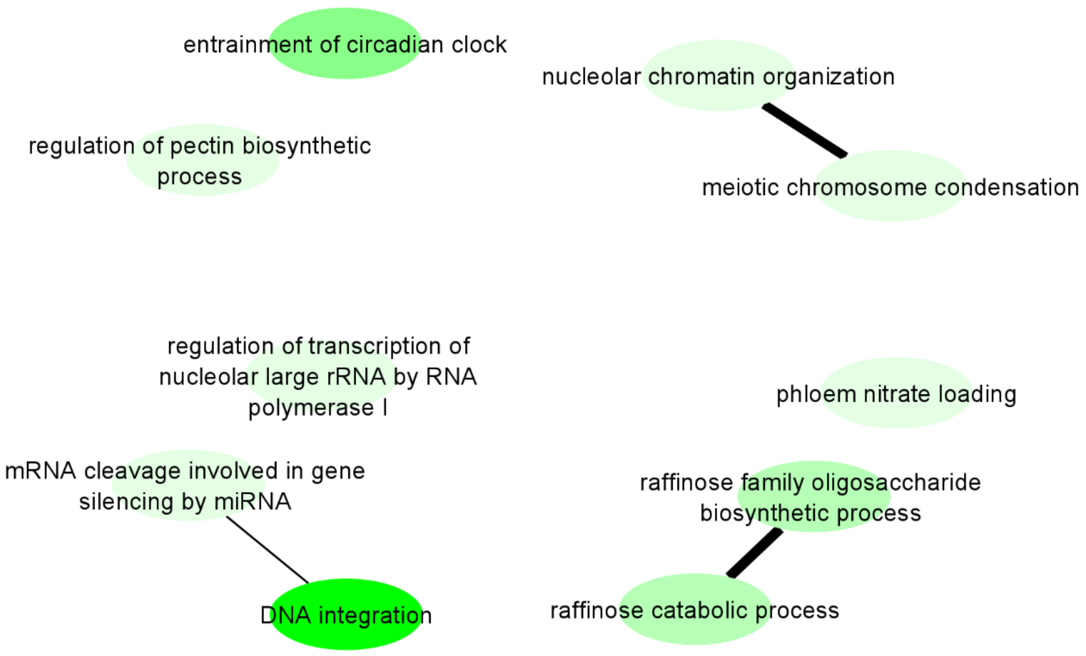


**Figure F10 –** Summary of biological processes enriched in the EQ Rapid suppression class under freezing temperatures (-4°C) in *Avicennia schaueriana*. The ellipses represent the enriched GO terms and the lines the genetic ontology relationship, the thicker the line the greater the ontogenetic relationship of the terms.

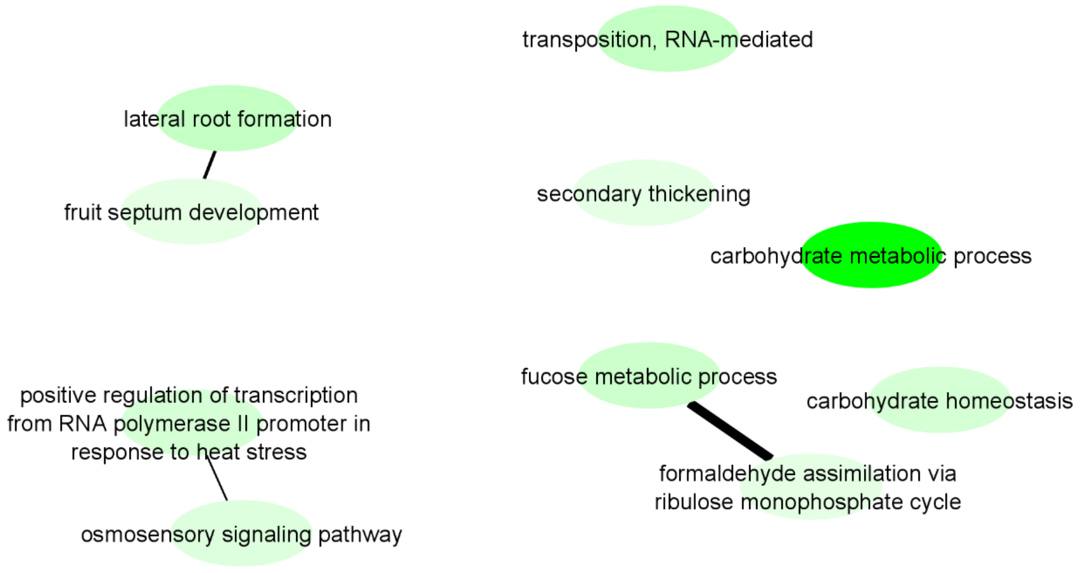


**Figure F11 –** Summary of biological processes enriched in ST Rapid suppression class in *Avicennia schaueriana* under freezing temperatures (-4°C). The ellipses represent the enriched GO terms and the lines the genetic ontology relationship, the thicker the line the greater the ontogenetic relationship of the terms.

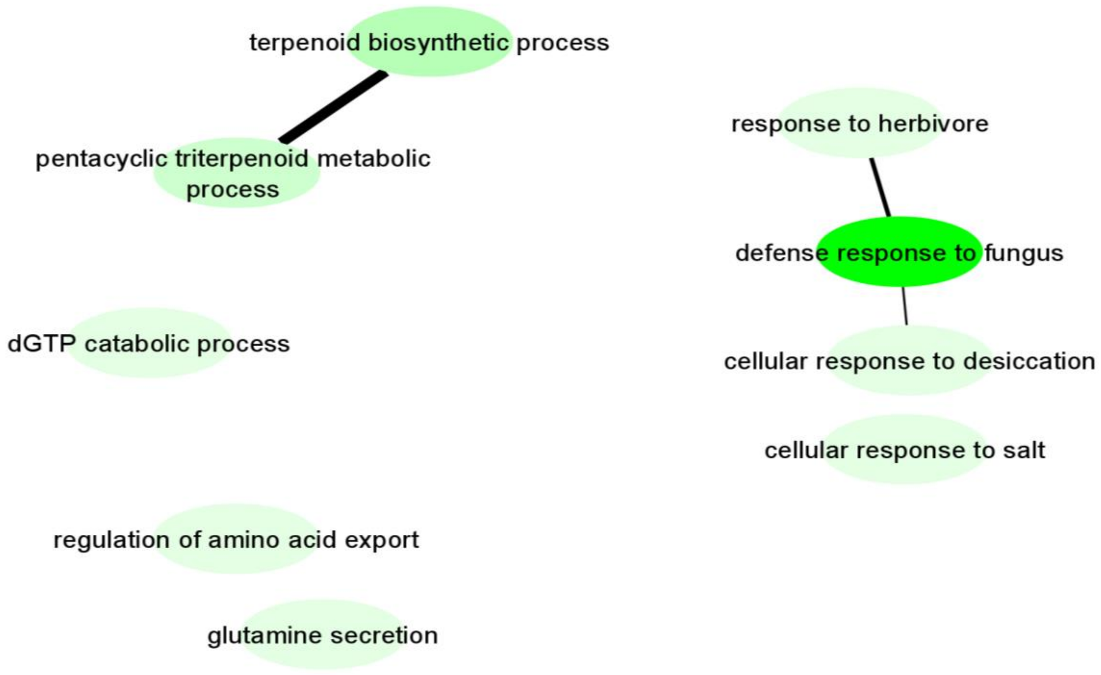


**Figure F12 –** Summary of biological processes enriched in the EQ Slow suppression class in *Avicennia schaueriana* under freezing temperatures (-4°C). The ellipses represent the enriched GO terms and the lines the genetic ontology relationship, the thicker the line the greater the ontogenetic relationship of the terms.

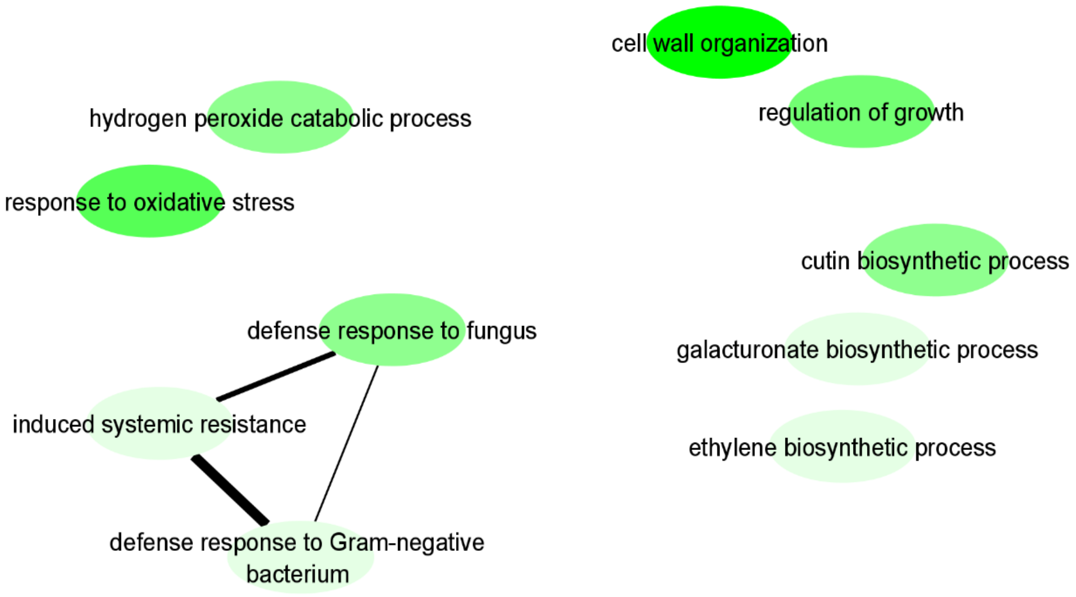


**Figure F13 –** Summary of biological processes enriched in the ST s Slow suppression class in *Avicennia schaueriana* under freezing temperatures (-4°C). The ellipses represent the enriched GO terms and the lines the genetic ontology relationship, the thicker the line the greater the ontogenetic relationship of the terms.

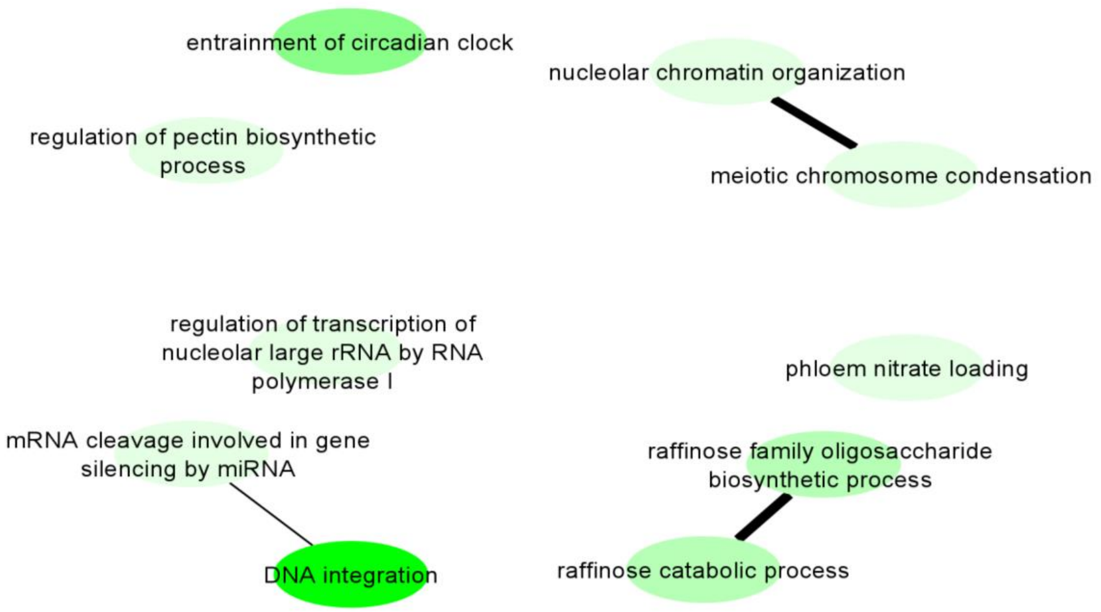


**Figure F14 –** Summary of biological processes enriched in the EQ Suppression followed by restitution class in *Avicennia schaueriana* under freezing temperatures (-4°C). The ellipses represent the enriched GO terms and the lines the genetic ontology relationship, the thicker the line the greater the ontogenetic relationship of the terms.

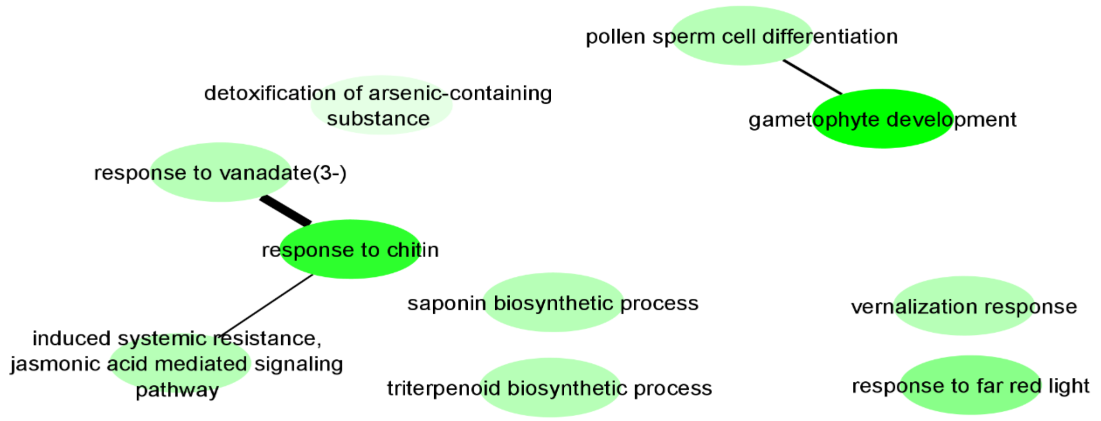


**Figure F15 –** Summary of biological processes enriched in the ST Suppression followed by restitution class in *Avicennia schaueriana* under freezing temperatures (-4°C). The ellipses represent the enriched GO terms and the lines the genetic ontology relationship, the thicker the line the greater the ontogenetic relationship of the terms.

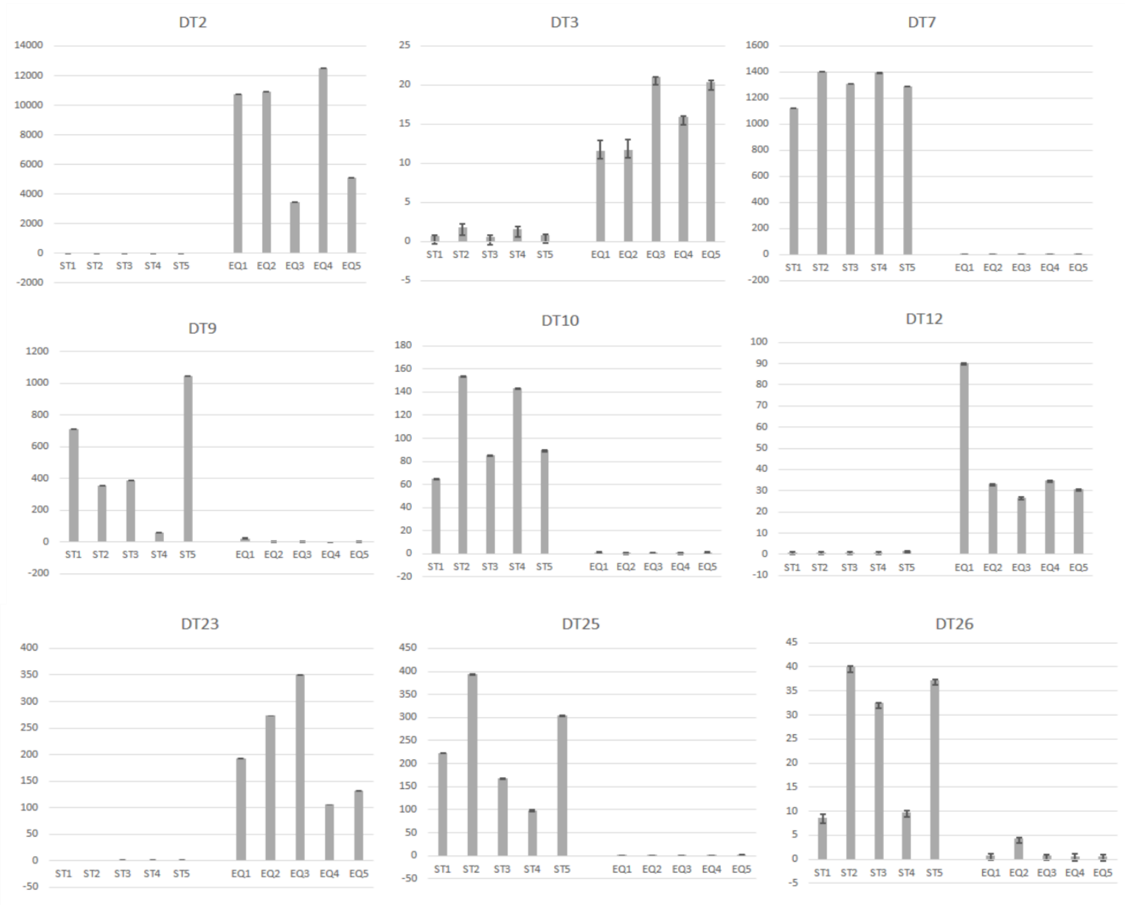


**Figure F16 –** Relative Quantification (RQ) of transcripts selected for validation via RT-qPCR. Relative expression per sample. The error bars correspond to the standard deviation between the technical triplicates of each reaction. All of these transcripts had a T test result < 0.05 when comparing the means of the two normalizing transcripts NR1 and NR2.
